# Evolution of the conformational ensemble and allosteric networks of apoptotic caspases in chordates

**DOI:** 10.1101/2025.02.14.638291

**Authors:** Isha Joglekar, Mithun Nag Karadi Giridhar, David A. Diaz, Ankit Deo, A. Clay Clark

## Abstract

Caspases, known for their role in apoptosis and inflammation, have recently emerged as potential regulators of cell differentiation and proliferation. The regulation of caspase conformations to affect these specific pathways is not well understood. Understanding the conformational landscapes, intermediate states, and mechanisms for fine-tuning these ensembles is critical for unraveling their diverse regulatory functions. We combined experimental and computational techniques to characterize the evolution of conformational ensemble in caspases. Using ancestral protein reconstruction, molecular dynamics simulations, and network analysis, we identify a network of residues that serve as a scaffold for conformational variation, and the network has been faithfully passed down over 500 million years of apoptotic caspase evolution. In addition, investigations of free energy landscapes provide thermodynamic insights into how initiator and effector caspase subfamilies stabilize monomeric and dimeric conformations while undergoing conformational dynamics. We find, through mutational in-vitro folding analysis, that the small subunit and amino acid networks at the base of the structure are essential for conformational dynamics beyond a ubiquitous intermediate present in the conformational ensemble of all apoptotic chordate caspases. We identify key allosteric hub residues that regulate the equilibrium in the conformational ensembles of all apoptotic caspases in chordates. Lastly, by combining our new data set with previous folding studies we provide a comprehensive model for folding of all apoptotic caspases in chordates.

## Introduction

Understanding how sequence variation in protein families leads to functional diversity and specificity while conserving the fold is a fundamental challenge in molecular evolution [1, 2]. Versatile functionalities in protein families arise from a complex and poorly understood choreography of conformational dynamics [3]. With the surge of data from sequence, structure, biochemical and biophysical studies, there is a need to integrate these diverse sources and understand their relationships to elucidate how protein families evolve amino acid interaction networks that drive dynamics while preserving the fold [4, 5]. Such efforts are important not only for advancing our understanding of molecular evolution, but also for developing more precise models of protein dynamics to inform pharmacological strategies by enhancing our understanding of the structure-function relationships in proteins [6].

Caspases are a class of enzymes that play a crucial function in the apoptosis and inflammation processes [7]. While apoptotic caspases have evolved into initiators, manifesting as stable monomers, and effectors, existing as stable dimers (Fig 1A), their activation mechanisms are paradoxically disparate [8]. Dimerization stands as a pivotal prerequisite for caspase activation, with initiator caspases tightly controlled through orchestrated dimerization within oligomerization platforms in response to apoptotic signals [7, 9]. Subsequent activation of initiator caspases triggers downstream activation of dimeric effector caspases, thus unraveling an intricate activation mechanism within the apoptotic cascade [10]. However, the phenomenon of dimerization is a multifaceted process that has not been thoroughly understood in either initiator or effector caspases [7, 11, 12]. Interestingly, investigations into the folding of the effector caspase subfamily have shed light on dimerization as a folding event, wherein partially folded, intermediate, conformations interact and fold into a dimeric state [13]. Further, dimerization has been reported to result in an average of a twofold increase in conformational free energy of effector caspases [14]. Moreover, the free energy of the monomeric conformation, which is the native state in initiator caspases, appears as a partially folded intermediate in the folding landscape of effector caspases. This suggests that the free energy of the monomeric conformation is conserved in both subfamilies [15]. Additional evidence supporting the monomeric conformation in effectors comes from kinetic folding data for procaspase-3, which indicates the presence of both dimerization-competent and dimerization-incompetent monomeric species [16]. This observation suggests that the subfamilies modulate the folding landscape to stabilize the monomeric conformation in initiators and guide effectors toward the dimeric state, leading to changes in regulation and function. The caspase family, with its well-characterized folding landscape coupled with numerous conformations in the structural database, offers an opportunity to explore how conformational stability is shaped by the evolution of amino acid networks and protein folding landscapes within protein families.

**Figure 1.**
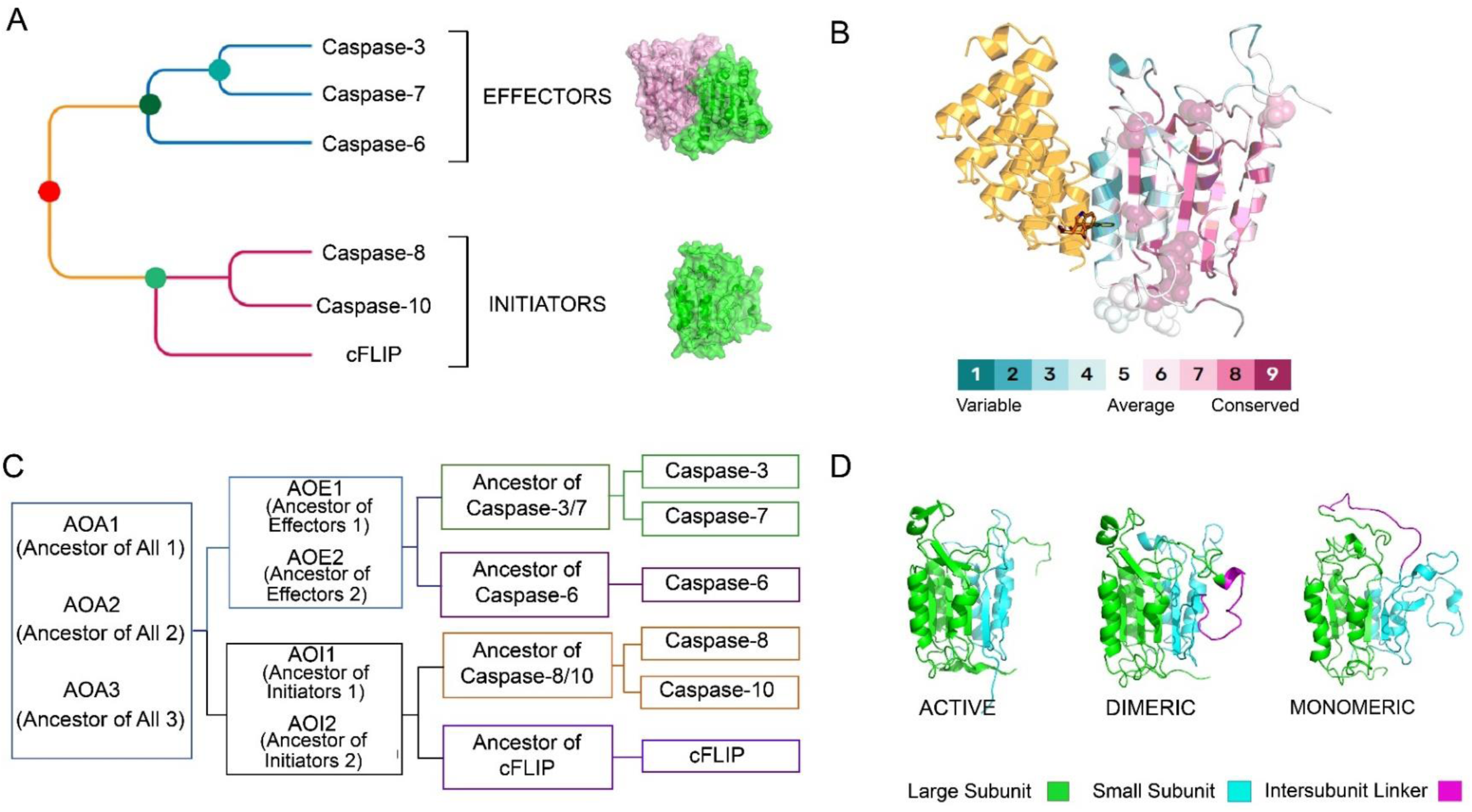
Caspase structure and evolution. (A) Evolutionary relationship of caspases involved in the extrinsic pathway of apoptosis. (B) Conservation of site-specific residue mapped on caspase-7, along with post-translational modification sites (depicted as spheres) and allosteric binding of DARPin to caspase-7. (C) Phylogenetic tree illustrating the evolutionary relationships among modern caspases and inferred ancestral nodes. (D) Structural representation of caspases in the active, dimeric and monomeric state, highlighting differences in their conformation, with the large subunit in green, small subunit in cyan and the inter-subunit linker in magenta.

In addition to sequence variations that influence the conformational energy landscapes, caspase subfamilies have evolved mechanisms to stabilize high energy states within the ensemble that play a role in non-apoptotic processes such as cell proliferation and differentiation [7]. Post-translational modification (PTM) of caspases is one such mechanism that affects caspase function by fine-tuning the ensemble of states in specific pathways [17]. Interestingly, conserved phosphorylation sites S347 in caspase-8, S150 in caspase-3, and T173 in caspase-7 are located in the same spatial position near α-helix 3 (Fig 1B) of the catalytic subunit and are highly conserved across organisms [18, 19]. These PTMs disrupt the active site loops and suggest a shared regulation model among diverse caspases [17]. A cluster of several PTMs, as shown in the same region (Fig1 B), depicts phosphorylation, nitrosylation, and ubiquitylation sites, near the conserved phosphorylation sites described above, making this region a hotspot for conformational regulation [18]. This provides an attractive avenue to explore conserved amino acid networks that govern conformational stability in caspases.

In order to characterize conserved amino acid networks and their evolution in the conformational landscapes of caspase sub-families, we employed ancestral protein reconstruction (APR), molecular dynamics (MD) simulations, essential dynamics (free energy landscape), network analysis, and mutational *in-vitro* folding studies [20–22]. Invertebrate sequences were removed from our APR because they are poorly described, and previous studies reveal that the folding patterns are under distinct evolutionary forces [8]. The presence of caspases-3,-6,-7, and -8-like homologs in non-chordates suggests that a pool of ancestral sequences acquired from the evolution of a common ancestor of Metazoa around 700 mya provided the framework for the evolution of their extant homologs in chordates [10, 22]. To simulate the ancestral pool, we reconstructed a pool of ancestral caspases (Fig 1C) named ancestor of all (AOA-1,-2,- 3) using chordate caspase sequences from three databases (Supplemental File 1). Furthermore, we reconstructed the ancestral sequences of all initiators (AOI-1,-2), ancestor of caspase-8/-10 (Anc8/10), and ancestor of cFLIP (Anc-cFLIP) (Figure 1C) from an additional initiator caspase database (Supplemental File 1). Ancestor of all effectors (AOE-1,-2), ancestor of caspase-3/-7 (Anc3/7) and ancestor of caspase-6 (Anc6), were reconstructed from previous databases compiled by Grinshpon et al [22]. (Figure 1C). A sequence alignment of all the reconstructed ancestral sequences with the extant human caspase sequences used in this study is shown in Supplemental Figure S1.

We performed molecular dynamics simulations in water and in urea with the active (intersubunit linker cleaved), dimeric (intersubunit linker intact), and monomeric (NMR solution structure of caspase-8: 2k7z) structures (Fig 1D) acquired from the PDB or modeled according to the nearest neighbor in the evolutionary tree (described in detail in the methods section). The NMR structure of caspase-8 (PDB ID: 2k7z) is the only available structure of a monomer; hence we utilized it to model all of the sequences [23]. Given the conserved free energy of the monomer in both subfamilies and the congruence of previous MD simulations with experimental findings, the 2k7z caspase-8 monomeric conformation stands as robust and ideal for modelling all the sequences [15].

Our data show that both monomeric and dimeric conformations exhibit an expansive ensemble of structures in contrast to the active conformation. MD simulations in water and in urea reveal that the small subunit has subfamily-specific dynamics that predate the caspase common ancestor in chordates. These studies provide insight into the energetic barriers that trap initiators in monomeric conformations while guiding effectors toward dimeric conformations as a result of amino acid interactions in the small subunit. Moreover, with in-depth network analysis of MD simulations, we systematically categorized stable scaffolding interactions within the hydrophobic core, and we show that the interactions have remained highly conserved since the common ancestor in all apoptotic caspases. In addition, we identify a network of residues in the small subunit with varied biochemical properties that are specific to the subfamilies. *In-vitro* folding analyses with the ancestral mutations swapped into subfamilies of extant caspases reveal the network to be critical to evolution of the protein fold and dynamics, beyond the identified stable core in apoptotic caspases. Limited trypsin proteolysis followed by mass spectrometry show that the mutations in the small subunit of the ancestral caspases affect the stability of an allosteric hotspot via a network of amino acid interactions in the loop regions at the base of the structure. Further we identify critical conserved residues in these networks that are located at the allosteric hotspot (Fig 1B) and that control conformational dynamics and fold beyond a conserved folding intermediate. The conserved amino acid network encompasses the stable conserved core in all apoptotic caspases. Lastly, we show that a network of conserved phenylalanine residues mediates allosteric signals to effect side chain packing in the hydrophobic core and hence affect global dynamics and caspase conformations.

## Results

### Conformational stability of apoptotic caspases in chordates is governed by varied packing of amino acid side chains between helices and beta sheets

We examined the stability of caspases using amino acid interaction networks [20]. In network analysis, amino acid residues are represented as nodes, and their interactions with neighboring residues are called edges [24]. Specifically, we utilized the degree centrality metric, which quantifies the number of non-covalent interactions that residues make with their neighbors [20]. The conformational dynamics of monomeric, dimeric, and active states are influenced by the arrangement and organization of residues in the core [25–27]. Utilizing degree metrics, we analyzed the spatial distribution of non-covalent bonds within distinct caspase conformations across the entire tree (Fig1C and 1D), providing insights into their evolution spanning a significant temporal scale of 500 million years [28].

The average degree data for each conformation, from molecular dynamics (MD) simulations in water, were parsed for each secondary structure element, and violin plots were utilized to display the distribution of these data for the active (Fig. 2A), dimeric (Fig. 2B), and monomeric (Fig. 2C) conformations. The violin plots for the monomeric conformation (Fig. 2C) show a considerable difference for the beta sheets and alpha helices compared to the dimeric conformation (Fig. 2B), indicating that the transition from monomer to dimer requires extensive rearrangements in these regions. In contrast, comparison of dimeric and active conformations shows smaller alterations, suggesting modest rearrangements in the transition from dimeric to active. Furthermore, subclassification of the degree scores into three categories based on Consurf conservation scores (Supplemental Figure S2) - high conservation (score of 9-8), intermediate conservation (score of 7-6) and variable (score below 5) - show that the conserved and intermediate conserved residues make the most contacts (non-covalent bonds) among the categories (Supplemental Figure S2 A,B,C) [29].

**Figure 2.**
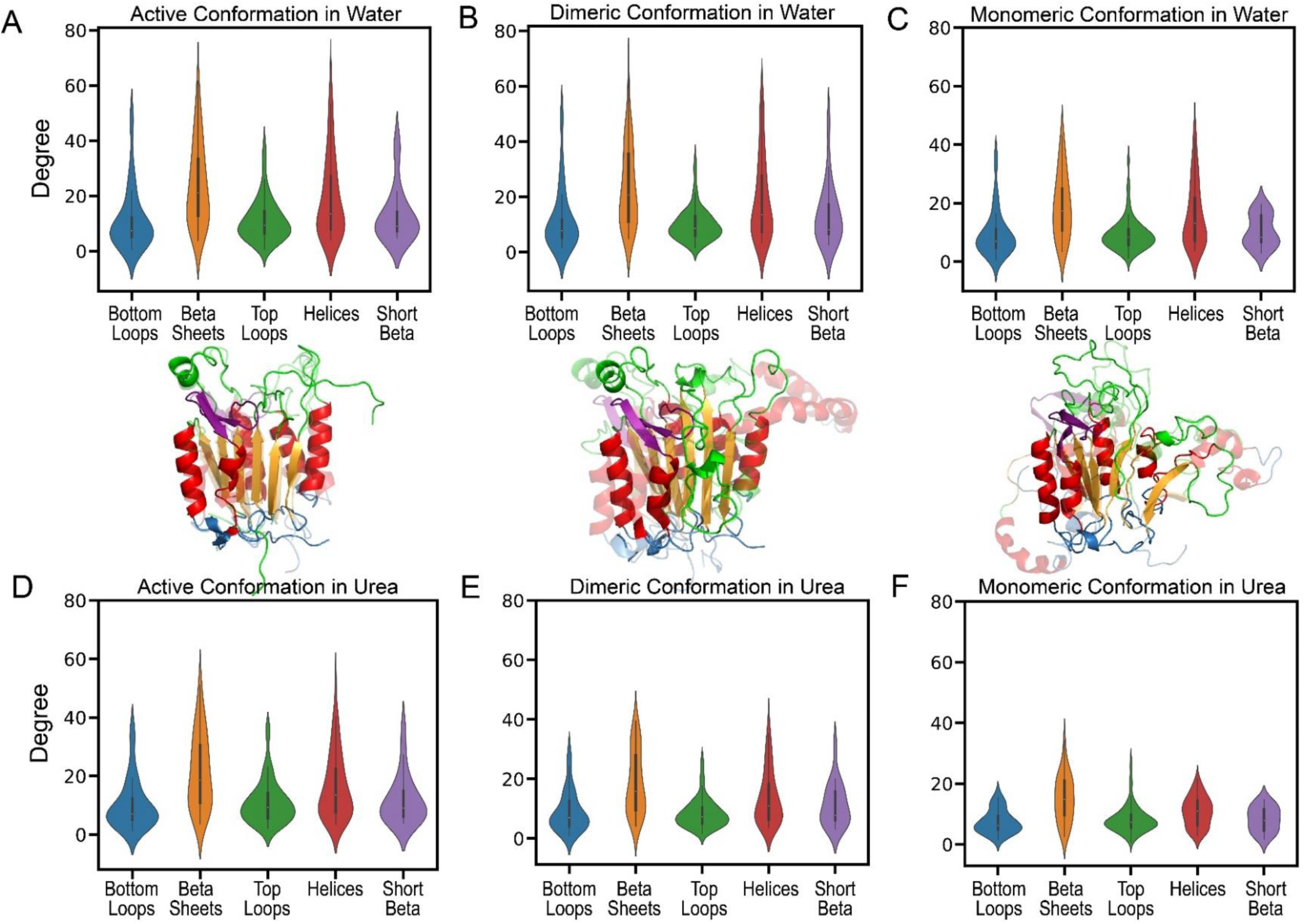
Violin plots representing the average degrees/contacts for the (A) active, (B) dimeric and (C) monomeric conformations in water and (D) active, (E) dimeric (F) monomeric conformations in 8M urea. Bottom loops (blue), beta-sheets (orange), catalytic site loops (green), helices (red) and the short beta-sheets (purple) are color-coded on the modeled caspase-8 structures. The solid structures represent the caspase-8 in water, and the translucent structures represent caspase-8 in 8M urea.

Similarly, we plotted degree data as violin plots for MD simulations in urea. The comparison of the violin plots for simulations in water (Figure 2A-C) to those in urea (Figure 2D-F) reveal that monomeric (Figure 2C,F) and dimeric (Figure 2B,E) conformations lose substantial non-covalent interactions, the majority of which are from loss of contacts in alpha helices and beta sheets. In addition, these plots show that the stability is mostly dictated by contacts in the beta sheets and helices that reside in the hydrophobic core. Moreover, sub classification suggests that conserved and intermediate-conserved residues display the maximum change for simulations in urea Supplemental Figure S2, panels D,E, and F, in comparison to those in water Supplemental Figure S2, panels A,B, and C.

Altogether, the data show that the stability of caspase conformations across a ∼500-million-year evolutionary span is significantly influenced by non-covalent interactions between beta sheets and alpha helices. Further, residues that make maximal contacts in the dimeric and active conformations are conserved throughout evolution. Notably, the monomeric conformation makes the fewest non-covalent bonds between helices and beta sheets, followed by a dramatic increase in the inactive dimeric conformation. Caspase dimerization is shown to contribute to ∼2-fold increase in the free energy of the monomer, and our results suggest that this increase is not simply a result of interface rearrangements in the small subunit to form the dimer but rather a function of global amino acid side chain rearrangements in the helices and beta sheets [13–15, 30]. The conformational free energy of the monomeric and dimeric inactive states correlates well with contact loss, offering insights into the high stability of the active conformation. We note that the conformational stability of the active dimer is unknown because folding is irreversible when the chain is cleaved.

### The small subunit displays conserved patterns of stability across initiator and effector subfamilies in monomeric and dimeric conformations

Since the active state is very stable and doesn’t display varied dynamics in our simulations, we characterized the evolution of the free energy landscape (FEL) of the dimeric and monomeric conformations of all the extant and ancestral caspases (Fig 1C) [31]. The essential dynamics were identified using Principal Component Analysis (PCA) based on molecular dynamics (MD) simulations in urea and in water [32]. The first two principal components (PCs), which explain much of the variance in the data, were utilized to generate a FEL for evolutionary analysis.

Comparing the FEL in urea (Fig 3A) to that of water (Supplemental Fig S3), a larger conformational space in urea is observed, indicating enhanced sampling of atoms. Despite simulation times of 200 ns in 8 M urea being insufficient for complete unfolding, these investigations can demonstrate relative stability across caspases, with a wider landscape indicating more unfolding. Moreover, the area or the width occupied by the FELs correlates with experimental findings. In the effector subfamily, the folding landscape of the dimeric conformation has been extensively studied [13, 14, 30]. Among the members of this subfamily, caspase-6 has been found to be the most stable when compared to caspase-3, caspase-7, and AOE-2. This observation is consistent with our findings, which indicate that caspase-6 exhibits the least accessible conformational space, as shown in Figure 3A, in comparison to caspase-3, caspase-7, and AOE-2. Currently, there are no folding data to quantify the conformational free energy of the dimeric conformation of caspases in the rest of the evolutionary tree, which limits our ability to provide a comprehensive evolutionary perspective. However, since our MD simulations in urea in the current and previous studies show excellent agreement with experimental findings, the data provide us an avenue to characterize the dimeric conformation in regard to an overall evolutionary perspective. Data from FEL plots (Fig 3A) suggest that the effector caspase lineage utilized the AOA-3 scaffold, with narrow FEL, from the ancestral pool, leading to a narrow FEL from AOE-1,-2 to the extant caspases-3,-6,-7, in essence stabilizing a dimeric scaffold in the conformational ensemble. In contrast, the initiator lineage evolved to exhibit a wider FEL from AOI-1,-2 to the extant caspases-8,-10, and cFLIP. Note that in the effector caspase lineage, only ancestor of caspase-3/7 shows a wide landscape, which is discussed later in this section.

**Figure 3.**
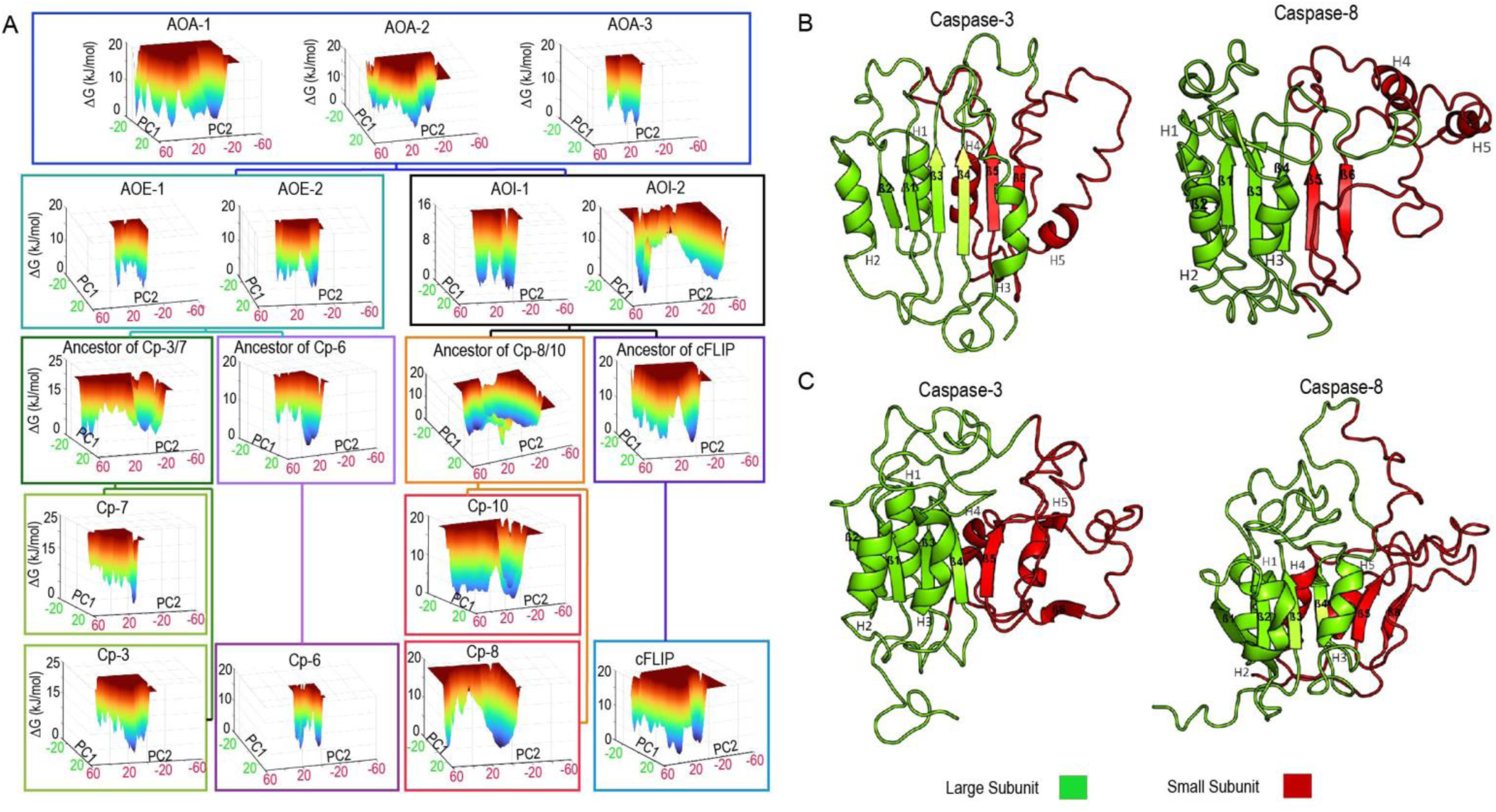
The free energy landscapes of the dimeric conformation of caspases in urea obtained from 200 ns MD simulations. (A) The FELs are generated as a function of projections of the MD trajectory onto the first (PC1) and the second (PC2) eigenvectors, respectively. Target FEL minima are represented as the metastable states visited during the simulations in urea for the dimeric conformation of (B) caspase-3 and caspase-8. Target FEL minima in the monomeric conformation for simulations in water are represented as the metastable states in (C) caspase-3 and caspase-8.

Given the insights garnered from FEL of simulations in urea, which indicate distinct stability profiles among caspase subfamilies in their dimeric conformation, we further analyzed the last metastable state that these systems visited to determine whether we could distinguish similar patterns of destabilization in each caspase subfamily [33]. In Figure 3 (panels B and C), we show examples metastable states of caspase-3 and of caspase-8 extracted from the FEL (Fig. 3A), which correspond to the last observed metastable state in urea for effector caspase-3 and initiator caspase-8, in dimeric conformation. The small subunit of caspase-3 (Fig. 3B) unfolds to a lesser extent than the small subunit of caspase-8 (Fig. 3C), and this trend is observed on average in the effector and initiator tree, respectively (Supplemental Fig S4). Hence, the broader landscapes observed in Fig. 3A for initiator caspases correspond to a less stable small subunit in the dimeric conformation. In caspase-3 (Fig. 3B and Supplemental Fig. S4), cFLIP, and other ancestors (Supplemental Fig. S4), helices 2 and 3 detach from the beta sheets. Note that the anti-parallel beta sheet 6 is destabilized in the ancestor of caspase-3/-7 (Supplemental Fig. S4), resulting in the destabilization of the small subunit, which gives rise to the larger FEL shown in Fig. 3A and is an exception in the effector caspase lineage. In the context of evolution and natural selection, it is conceivable that caspase-3/7-like sequences may have emerged within the ancestral gene pool through gene duplication without adversely affecting the fitness of the ancestral organism. This could have subsequently paved the way for neofunctionalization in caspase-3 and caspase-7. These results highlight the trial-and-error nature inherent in evolution and natural selection, shaping the design of specialized enzymes across extant organisms.

For the monomeric conformation in urea, there is no discernible difference in the FEL between initiators and effectors (Supplemental Fig. S5). However, in these simulations with the same urea concentration (8M), the FEL for the monomeric conformation (Fig. S5) is broader (PC1) than the dimeric conformation (Fig. 3A), indicating extensive unfolding of the less stable monomeric conformation. Interestingly, for simulations in water, the average MD conformation reveals that beta sheet 6 unfolds in caspase-3, but the monomeric conformation of caspase-8 is relatively stable (Fig. 3C) which is observed to be an average trend in the effector and initiator subfamilies (Supplemental Fig. S6), respectively.

In summary, conformational landscapes of monomeric and dimeric caspases were already established in the common chordate ancestor, and their variable stability has been passed on faithfully in each subfamily. Simulations, in urea, of the entire caspase tree suggest that the dimeric conformation of the initiator caspase lineage is less stable compared to the dimeric conformation of the effector caspase lineage. The data suggest that the lower stability is due to weakened interactions between helices 4 and 5, and beta sheets 5 and 6. Conversely, simulations of caspases in water reveal that the effector caspase lineage is unstable in the monomeric conformation, particularly due to destabilizing interactions near beta sheet 6. Overall, these studies indicate that amino acid interactions in the small subunit, and their changes throughout evolution, are largely responsible for changes in the conformational landscape of the initiator and effector subfamilies.

### In caspase sub-families, the small subunit and co-evolving networks of residues evolve folding landscapes while also demonstrating an implicit connection to an allosteric hotspot

We identified residues within the small subunit that display distinct biochemical properties between the initiator and effector subfamilies in the entire chordate database (Supplemental File S1). These 21 residues (Fig. 4A) map to the small subunit, as depicted by the caspase-8 structure in Figure 4A and the alignment in Supplemental Figure S1. We hypothesized that these 21 residues influence the observed conformational diversity among caspase subfamilies. We reasoned that by exchanging the 21 mutations in caspase-3 and caspase-8, it might be possible to influence the monomer-dimer equilibrium in caspases-8 and -3.

**Figure 4.**
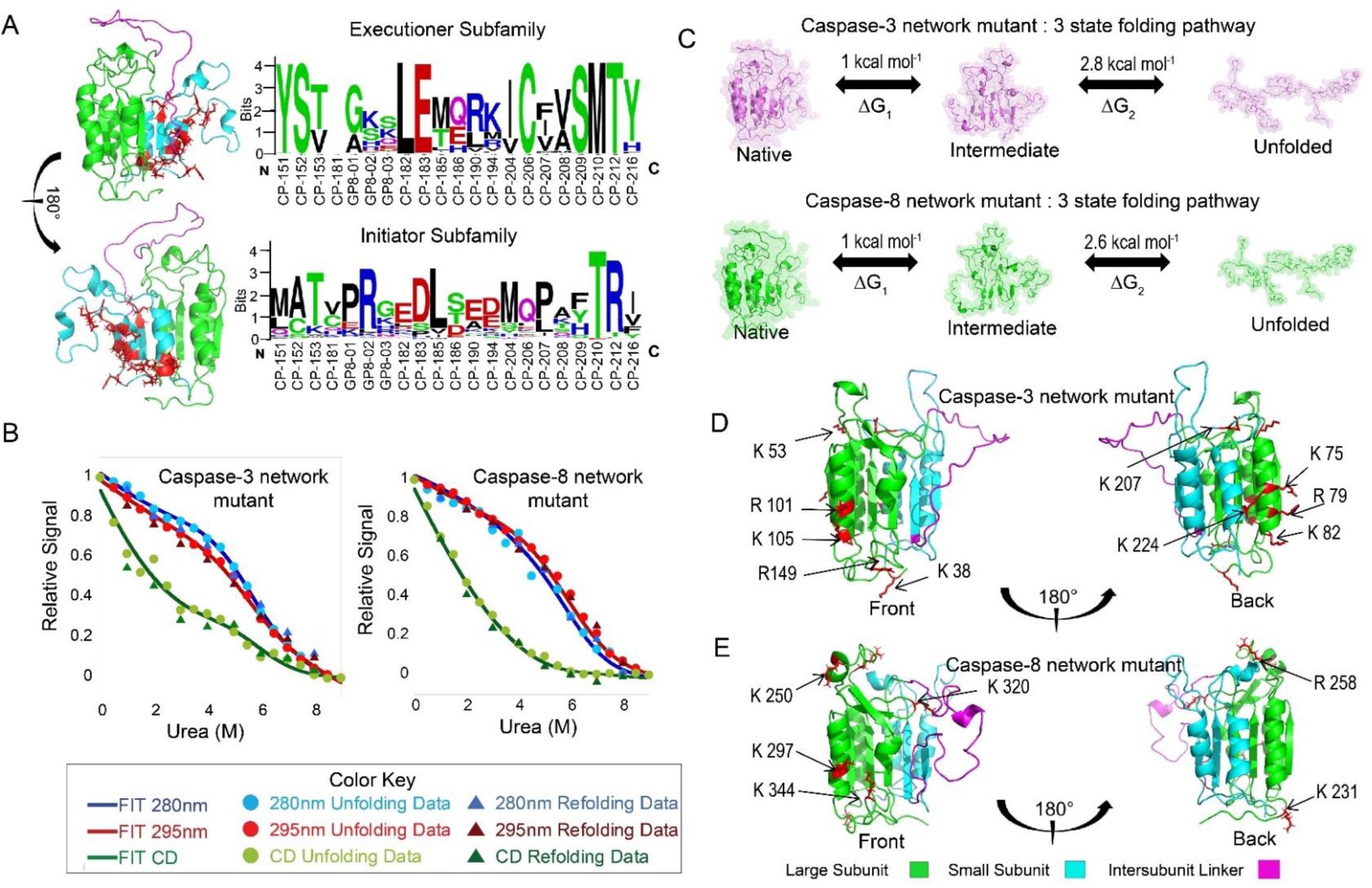
(A) Site-specific amino acid differences within subfamilies in the small subunit represented on a web logo and mapped onto the caspase-8 monomer. The large subunit is depicted in green, the small subunit in cyan, the inter-subunit linker in magenta and site-specific differences are mapped as red sticks onto the structure. (B) Equilibrium unfolding of caspase-3 network mutant and caspase-8 network mutant at pH 7.5 monitored by fluorescence emission with excitation at 280 nm ( ), 295 nm (. ), CD ( ) and refolding at 280 nm ( ), 295 nm ( ) and CD ( ). As described in the text, solid lines represent global fits to the data at 280 nm ( ), 295 nm ( ) and CD ( ). (C) Representation of the free energies obtained from the global fits for each step of unfolding for caspase-3 network mutant and caspase-8 network mutant. Cleavage products of limited trypsin proteolysis at pH 7.5 were determined by MALDI-TOF MS as described in the text and represented on caspase-3 network mutant structure (D) and caspase-8 network mutant structure (E). The large subunit is depicted in green, the small subunit in cyan, inter-subunit linker in magenta and the cleavage sites as red sticks.

We undertook *in-vitro* folding investigations with these 21 mutations (21M) in caspase-8 and caspase-3 to test the veracity of our idea. We tracked urea-induced unfolding and refolding of tertiary structure using excitation wavelengths of 280 nm (all aromatic residues) or 295 nm (tryptophan residues only) and fluorescence emission from 300 nm to 400 nm. We also performed far-UV circular dichroism for 21M caspase-3 (Supplemental Fig. S7 A,B,C) and 21M caspase-8 (Supplemental Fig. S7 D,E,F) to monitor changes in secondary structure. Our folding studies revealed no dependence on the protein concentration for caspase-3 21M (Supplemental Fig. S7G) or for caspase-8 21M (Supplemental Fig. S7H). The findings suggest that the equilibrium of monomer-to-dimer in caspase-3 was shifted to the monomeric state. However, the comparable mutations did not result in a stable dimer for caspase-8. Unexpectedly, the folding data for caspase-3 21M and caspase-8 21M show that the folding patterns and thermodynamic parameters exhibit a remarkable similarity (Fig. 4B), and data sets for both mutants can be characterized by a three-state folding model for a monomer (Fig. 4C), in which the native monomer unfolds to a partially folded intermediate prior to unfolding (N ν I ν U). For both proteins, the mutations result in substantial changes in stability compared to their wild type counterparts [14, 15]. The thermodynamic parameters extracted from the fits of the data (Table S1), show that the first transition (N ν I) exhibits a conformational free energy of ∼1 kcal/mol, while the second transition (I ν U) exhibits a conformational free energy of ∼2.7 kcal/mol. Thus, both caspases with 21 mutations in the small subunit exhibit a total conformational free energy of ∼3.7 kcal/mol.

We reasoned that the mutations destabilize the small subunit due to interrupting the networks of coevolving residues in the large subunit. Consequently, these 21M proteins became confined to an evolutionarily conserved, partially stable, folding intermediate. Overall, the mutations appear to render the proteins incapable of accessing other conformations within their respective ensembles, as opposed to their wild-type counterparts which fold into a stable dimer (caspase-3) or monomer (caspase-8). To further identify regions that were destabilized because of the 21M mutations, we performed limited trypsin proteolysis and used MALDI-TOF mass spectrometry to identify regions of the proteins with the highest cleavages at a time point of one hour. The results were compared for wild-type caspase-3 (Supplemental Fig. S8A), caspase-3 21M (Supplemental Fig. S8B), and caspase-8 21M (Supplemental Fig. S8C). Notably, identical conditions to those described in our previous report were used for caspase-8 wild-type; consequently, we compared the results of the 21M mutants to wild-type from our previous investigations for caspase-8 [15]. We compared the top 10 cleavages for caspase-3 wild-type (Supplemental Fig. S8D) and caspase-3 21M (Supplemental Fig. S8E) and for caspase-8 wild-type (Supplemental Fig. S8F) and caspase-8 21M (Supplemental Fig. S8G) to determine regions that are destabilized in their 21M versions. Overall, the data for caspases-3 and -8 (Figures 4D and 4E, respectively), indicate that helices 2 and 3 and loop regions in the large subunit that are otherwise inaccessible to the protease in the wild-type proteins are destabilized in the 21M variants. These findings indicate that mutants in the small subunit affect the stability of helices 2 and 3 in the large subunit, with a particular clustering near an allosteric hotspot. The results suggest that interactions in co-evolving networks of amino acids link the small subunit to the allosteric site.

### Highly conserved residues that exhibit high degree centrality and betweenness centrality provide a scaffold for the evolution of conformational dynamics in the subfamilies

In order to identify co-evolving networks and explore the inherent link between the small subunit and the allosteric pocket on the large subunit, we first classified conserved networks in the central core as well as other evolving networks in the protein sequences. We employed degree centrality (DC) and betweenness centrality (BC) metrics to discern critical elements in the networks [20]. These metrics allowed us to identify residues with numerous interactions and to distinguish interactions that may be central to communication pathways within the protein structure.

To identify nodes in identical spatial position that display high values for these metrics across evolutionary time, we averaged positional values across the entire tree, for both the monomeric and the dimeric conformations, separately. The active conformation was excluded from the analysis due to its inherent stability and limited dynamical behavior. Pair plots of degree vs betweenness centrality for the average in the dimeric (Supplemental Fig. S9A) and monomeric (Supplemental Fig. S9B) conformations show that most values cluster below 20 for degree and 1000 for betweenness. Residues that fall outside this quadrant in both the monomeric and dimeric conformations represent stable scaffolding interactions around which conformations fluctuate [34]. We mapped high DC and BC residues in all caspases onto average weighted network maps obtained from molecular dynamics (MD) simulations for both monomeric and dimeric conformations of caspase-3 (Supplemental Fig. S9C and S9D) and of caspase-8 (Supplemental Fig. S9E and S9F). The data demonstrate pronounced thick edges, extensive interconnections, and central role for each high DC and BC amino acid within the entire network of amino acid interactions. Subsequently, the residues were mapped onto the structure of caspase-8 (Fig. 5A), and predominantly occupy positions within the hydrophobic core. The results highlight the integral role for these residues in facilitating information transfer throughout the entire molecular structure. These residues belong to the intermediate and well conserved groups (Supplemental Fig. S1A and S1B) in the Consurf grouping described above. The intermediate-well conserved residues predominantly populate the region of the molecule near helices 2 and 3. The less selective evolutionary pressure on these residues likely results in variable stability of helix 2 and 3 observed in some caspases in our MD simulations (Supplemental Fig. S4). On the contrary, high DC and BC residues on the opposite side of the structure are highly conserved. Interestingly, in MD simulations of unfolding in urea (Supplemental Video 1), observed in reverse, the highly conserved residues are the first to collapse or fold, followed by the intermediate-conserved residues. As conformations vary beyond the monomer, the highly conserved residues exhibit minimal dynamics, followed by the intermediate-well conserved, and then the other elements around them are prone to maximal dynamics.

**Figure 5.**
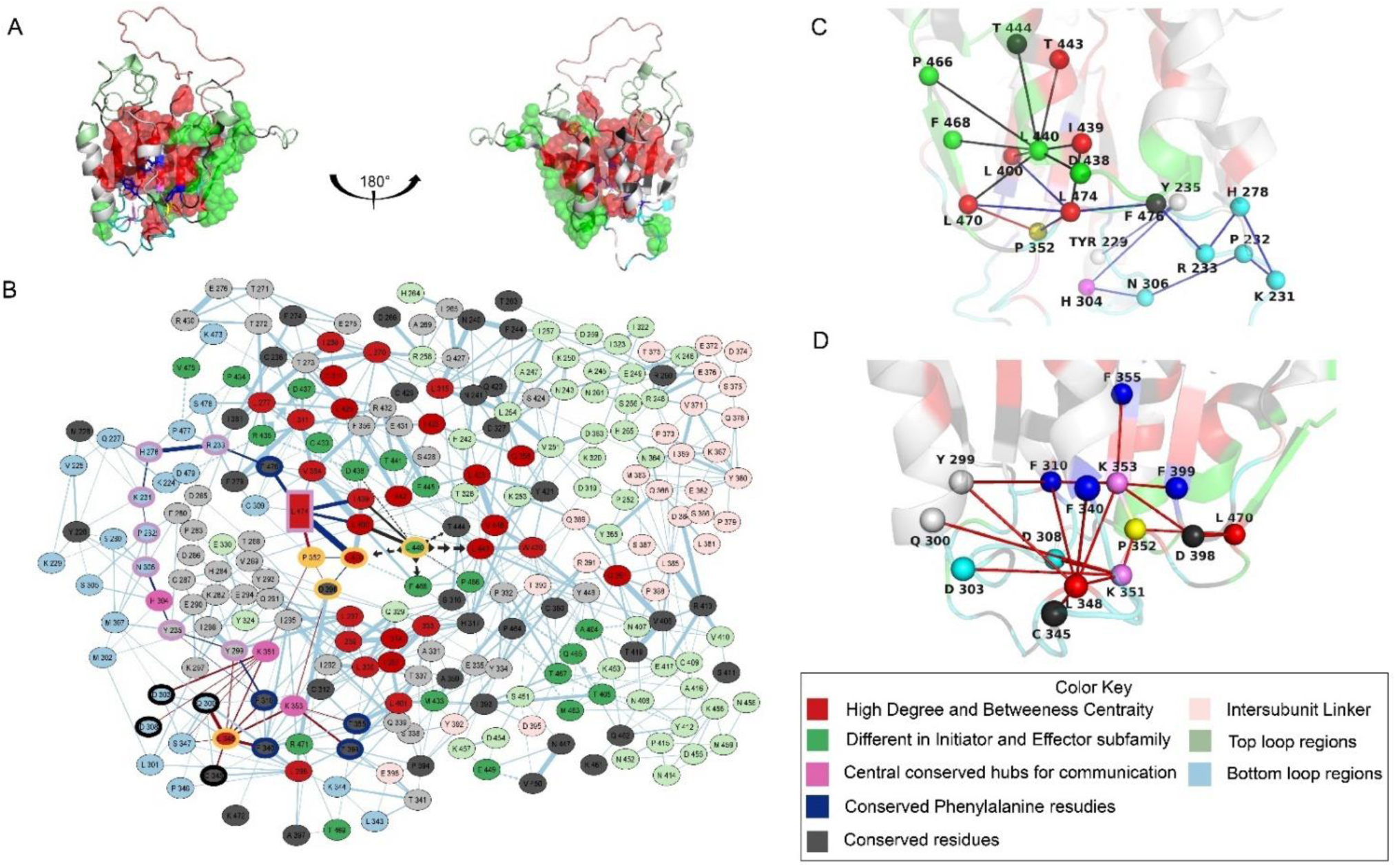
(A) MD average structure of monomeric caspase-8 in water highlighting residues having a high degree and betweenness centrality as red spheres, residues that are different within the initiator and effector subfamilies as green spheres and the conserved phenylalanine residues in the core as blue sticks. (B) A 2D representation of the network map of monomeric caspase-8 in water highlighting conserved residues as grey nodes, residues on the inter-subunit linker as light pink nodes, residues on the top loops as light green nodes and residues in the bottom loops as light blue nodes. Interaction networks mapped onto the caspase-8 monomeric structure for residues on the bottom loops at the back (below helices 1 and 4) (C), and residues on the bottom loops on the front (below helices 2 and 3) (D).

The 21M mutations in the small subunit (Fig. 4A) are absent from the high DC and BC set of residues (Fig. 5A and Supplemental Fig. S1), indicating they are dynamic as conformations vary. In the network of the 21 mutations in caspase-8, L440 (at the bottom of helices 4 and 5) acts as a central hub (Fig. 5, panels B and C), making the most stabilizing contacts at the bottom loops, and is highly conserved in all initiator caspases (Fig. 4A and Supplemental Fig. S1). Moreover, research findings indicating that interactions within the bottom loop region contribute significantly to dimer stability, accounting for 30-50% of dimer stability [35]. The shortest pathway for mutations around the L440 hub to affect the allosteric hotspot is facilitated by a critical residue, P352, located on β-sheet 4, which is also the location of the catalytic cysteine (Fig. 5B and 5C). Proline can adopt cis and trans isomers, and the highly conserved P352 is at a critical location that links high DC and BC networks located on one side of the protein to those on the other side. Further, P352 is covalently connected to highly conserved K351 and K353, both of which bury into the hydrophobic core at the front face of the protein (Fig. 5B and 5D). Hence, changes to the interaction network of P352 impacts the packing of residues on the opposite face of the protein. This phenomenon is exemplified in morph videos featuring the monomeric and dimeric conformations of caspase-8 (Supplemental Video 2). Notably, as conserved tyrosine Y226 (N-terminus) packs into the bottom loop region in the rear, P352 is altered, impacting the arrangement of high DC and BC residues at the front (Supplemental Video 2). Thus, the data show that an alternate route to destabilize helices 2 and 3, and subsequently the allosteric pocket, involves a distinct path that includes changes to positioning of conserved K351, K353 and P352 (Fig 5B and 5D).

In summary, by employing degree and betweenness centrality measures, we identified a conserved network of non-covalent interactions between β-sheets and α- helices (Fig. 1 and Supplemental Fig. 2), which are central to stability and communication. Given that the folding and stability patterns of caspase-3 21M and caspase-8 21M converge (Fig. 4B & C), this convergence may be a result of conserved elements spanning over 500 million years of evolution, corresponding to the identified high DC and BC residues (Fig. 5A). Our network analysis suggests that the 21 mutations in the small subunit ultimately destabilize co-evolving networks at the base of the structure that in turn affect the allosteric pocket through destabilizing helices 2 and 3 of the opposite face of the protein. Further, subtle modification of core residues around helices 2 and 3 allows for varied allosteric regulatory mechanisms in apoptotic caspases. This evolving allosteric network suggests differential modulation of catalysis in apoptotic caspases by affecting the critical histidine above helix 3 and dimerization via the small subunit in ways unique to each caspase.

### Conserved residue hubs at the base of the structure control the active site cysteine and histidine dyad while also regulating conformational dynamics

Conformational alterations driven by global shifts in residue interaction networks are frequently modulated by critical hub residues that govern conformational landscapes and allosteric signaling in proteins [36–38]. Here, we observed that all of the communication signals from the small subunit that alter helices 2 and 3 traverse bottom loop regions and appear to mainly modulate conserved residues H304, K351 and K353. The modulations create global rearrangements in the protein structure and may be involved in trapping the conformational ensemble of the 21M mutants in the partially folded state described above. To examine these residues further, we performed alanine scanning mutations of these residues in caspase-8 and characterized changes in the folding landscape.

Similar to the 21M mutants (Fig. 4B), described above, we examined urea-induced unfolding and refolding using fluorescence emission and circular dichroism spectroscopies for caspase-8 H304A (Supplemental Fig. S10 A,B and C) and caspase-8 K351A + K353A (Supplemental Fig. S10 D,E and F). The results show that the folding model shifts to a two-state pathway for caspase-8 H304A and K351A + K353A mutants (Fig. 6A and 6B) suggesting that the native state observed in caspases-3 and -8 21M (Fig. 4C) is destabilized in favor of the partially folded intermediate. The folding trends (Fig. 6A) and the parameters (Supplemental Table S1) show that the conformational free energy of intermediate seen in the 21M mutant (Fig 4C) is not affected. Altogether, the data suggest that replacing K351, K353, and H304 with alanine destabilize caspase-8 such that the “native” conformation is similar to the partially folded intermediate (I) observed in the 21M variants.

**Figure 6.**
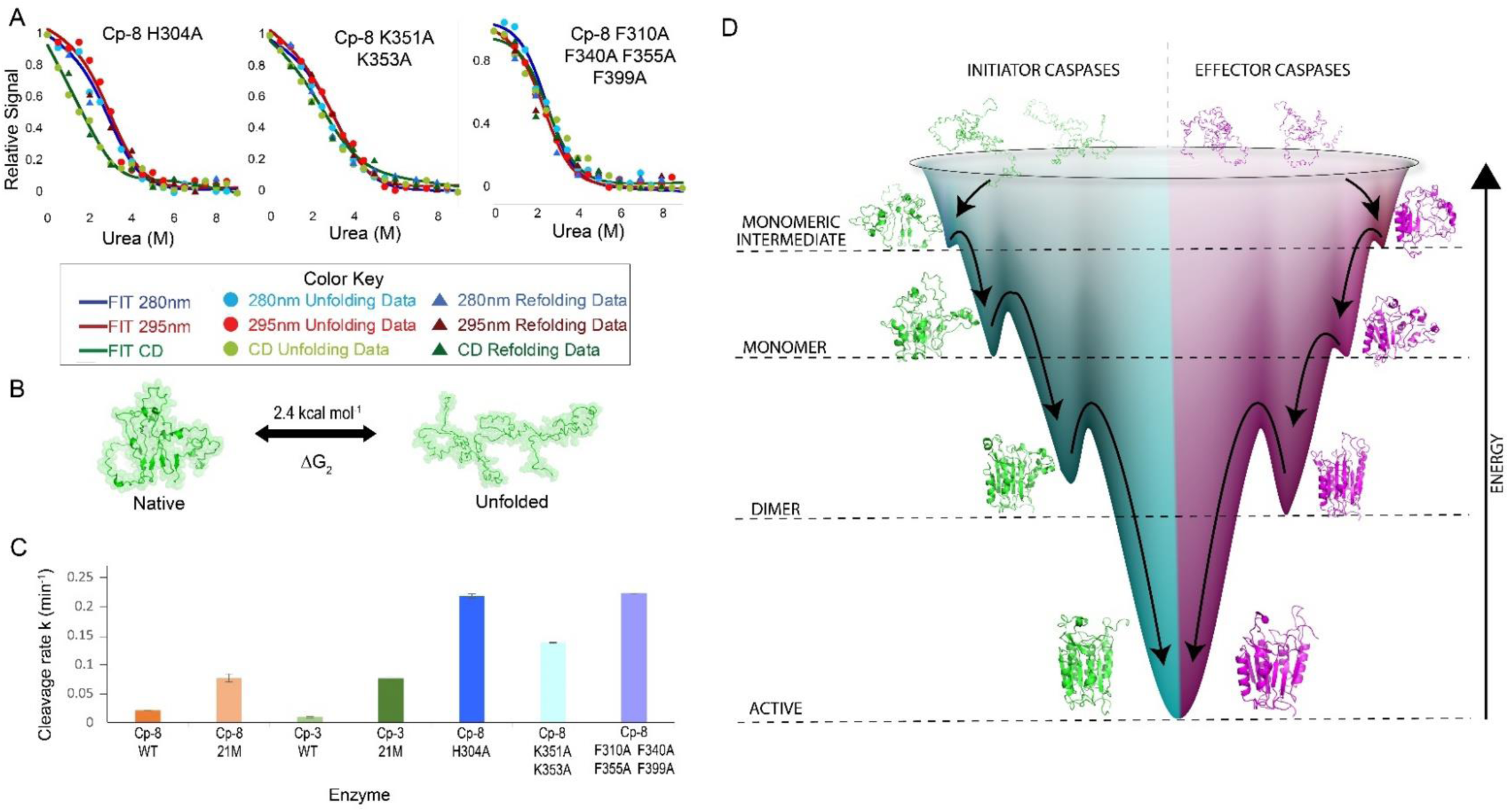
(A) Equilibrium unfolding of caspase-8 H304A, caspase-8 K351A + K353A and caspase-8 F310A + F340A + F355A + F399A at pH 7.5 monitored by fluorescence emission with excitation at 280 nm (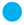), 295 nm (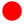), CD (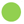) and refolding at 280 nm (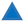), 295 nm (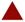) and CD (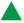). As described in the text, solid lines represent global fits to the data at 280 nm (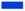), 295 nm (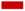) and CD (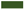). (B) Representation of a two-state unfolding model and free energy obtained from the global fits of unfolding for caspase-8 mutants. (C) Kinetics of cleavage of annotated bands from limited trypsin proteolysis (Supplemental Figures S8 and S11) were to fit to single exponential equation to determine the apparent rate constant, as described in the text. (D) A folding funnel model for the conformational stability of caspases.

After classifying high DC and BC residues, we identified highly conserved F310, F340, F355, and F399 that, despite being in the hydrophobic core near the allosteric pocket (Fig. 5D), do not appear among the high DC and BC residues, indicating their dynamic behavior as conformations vary from monomer to dimer. As the protein structure progresses from monomer to dimer to active conformations, these highly conserved phenylalanine residues appear to transmit changes at the bottom loop regions to the active site by rearranging the high DC and BC side chains in the core (Supplemental Video 3). Hence, we hypothesized that mutating these residues would not affect the stability of the partially folded intermediate observed in the 21M mutants (Fig. 4C), or as we speculate, the comparable “native” state caspase-8 H304A and K351A +K353A variants (Fig. 6A). We mutated the four phenylalanine residues to alanine and determined changes in the folding and stability of the protein (Supplemental Fig. S10 G, H, and I). Surprisingly, caspase-8(F310A,F355A,F355A,F399A), called caspase-8(F/A), exhibited an increase in secondary structure compared to the H304A or K351A/K353A variants (Supplemental Fig. S10 C,E and F). For the protein with the F- to-A mutations, the data are well-described by a two-state folding model (N ν U) (Fig. 6A and 6B) with a conformational free energy of 2.3 kcal/mol (Supplemental Table S1), which is identical to those of the H304A and K351A/K353A caspase-8 mutants (Fig. 6A). Together, the results agree with our hypothesis, suggesting that the four phenylalanine residues have similar effects to the native structure as H304, K351, and K353 by destabilizing the native conformation relative to the partially folded intermediate. However, since the residues have fewer contacts, as determined by lower DC and BC values, their role likely involves varying conformational stability through modulating the packing of side chains in the hydrophobic core.

We examined the caspase-8 variants by limited trypsin proteolysis coupled with mass spectrometry, and we show that the allosteric pocket is destabilized in caspase-8 H304A (Supplemental Fig. S11 A and B), K351A + K353A (Supplemental Fig. S11 E and F) and F310A + F340A +F355A + F399A (Supplemental Fig. S11 E and F). Further, the rate of cleavage of the native protein band correlates with our folding models. For example, caspase-8 21M and caspase-3 21M, both of which unfold via a three-state pathway, are cleaved at a similar rate, albeit at a higher rate than their wild-type counterparts. The data also show that caspase-8 H304A, caspase-8(K351A + K353A) and caspase-8(F/A), which unfold via a two-state pathway, are cleaved at a higher rate than the 21M variants (Fig. 6C). Overall, the data correlate well with our conclusion that mutations affecting the packing of the hydrophobic core or the conserved interaction network destabilize the native conformation of the monomer such that the protein is partially folded.

Overall, our studies suggest that the high DC and BC residues collapse early in refolding and substantially contribute to the stability of the partially folded intermediate in the folding transition of U to I (Fig. 4C). In addition, the high DC and BC residues have been conserved for more than 500 million years, with little modification in the intermediate-conserved region near the allosteric pocket (Supplemental Fig. S1 A and B). In caspase-8, H304, K351 and K353 are part of a conserved interaction hub, and alterations around the hub are sensed by several phenylalanine residues (F310, F340, F355 and F399), which modulate packing in the hydrophobic core. We analyzed the crystal structures for the ancestor of effector caspases [22] and the ancestor of caspase-6 [22] with eMap [39], and the results suggest that the highly conserved phenylalanine and cysteine residues in the core serve as electron or hole hopping channels in caspases to transport charges rapidly over long distances (Supplementary Fig. S12) [39, 40]. We suggest that caspases might use these transfer channels to modulate enzyme activity through changes in conformational dynamics. Lastly, K351, P352, K353 and F355 are located on β-sheet 4, which carries the catalytic cysteine, hence making this strand a vital element in catalysis and conformational regulation. The highly conserved H304, which modulates helices 2 and 3 in the allosteric pocket, plays a crucial role in regulating the position of the catalytic histidine. Collectively, our study provides mechanistic insights into allosteric control mechanisms that influence catalysis and regulate the conformational diversity within apoptotic caspases.

## Discussion

Caspases are essential enzymes in the regulation of apoptosis and the inflammatory response; however, the involvement of caspases in additional forms of cell death and non-cell death processes, such as differentiation and proliferation, remains poorly known [41]. Understanding caspase conformational landscapes, partially folded intermediate states, and mechanisms to fine-tune the conformational ensemble is crucial for comprehending regulation and, consequently, the fine-tuning of conformation to affect specific pathways [7]. In this study, we used ancestral protein reconstruction and MD simulations to identify a conserved network of residues that provide a scaffold for the evolution of subfamilies and their conformational dynamics, and with network analysis and FEL, we demonstrate how initiators and effectors stabilized the monomeric and dimeric folds through their changes in conformational dynamics over evolutionary time.

The folding funnel depicted in Figure 6D highlights our findings in conjunction with comprehensive experimental folding research conducted previously for initiator and effector caspase subfamilies [13–15, 30]. Network analysis for simulations in water and in urea demonstrate that the active conformation is the most stable, followed by the dimeric and monomeric conformations (Fig. 1), which is illustrated as energy gradients for these conformations in the folding funnel (Fig. 6D). The active conformation depicts the lowest minima in the folding funnel (Fig. 6D) and indicates maximum stability among all conformations. There are no experimental conformational free energy data for initiator caspases in the dimeric conformation; however, insights gleaned from our FEL analysis (Fig. 3A) show that the dimeric conformation is less stable in initiator compared to effector caspases, resulting in a comparably lower minimum for initiator caspases in the folding funnel (Fig. 6D). In the monomeric conformation, our data suggest that effector caspases are less stable than initiator caspases due to a spontaneous loss of high-degree interactions between the anti-parallel β-sheet and the core of the molecule, as observed in MD simulations in water (Fig. 3C and Supplemental Fig. S4), which has been represented by a higher energy barrier for the initiator caspases in the folding funnel model (Fig. 6D).

We identified 21 residues in the small subunit that are evolving distinct biochemical properties within each subfamily (Fig. 4A). *In-vitro* folding experiments with these residues swapped between caspase-3 and caspase-8 indicate that the otherwise distinct folding patterns of the wild-type proteins converge to a similar 3-state folding pathway in these mutants (Fig. 4C) [13, 15]. The 21 mutations in the small subunit disrupt the native states of caspase-8 and of caspase-3, stabilizing a partially folded conformation with a free energy of ∼3.7 kcal mol^-1^. The data suggest that the mutations perturb co-evolving networks coupled to an allosteric pocket (Fig. 4D and 4E). The free energy of the intermediate conformation reported in this 3-state pathway is 2.7 kcal mol^- 1^, which is identical to the free energy of the intermediate observed in the wild-type caspase-8 folding landscape [15]. The presence of the intermediate within the folding landscapes of evolutionarily distant caspase-3 and caspase-8 strongly implies its ubiquity among all chordate caspases. Supporting evidence from evolutionary network analysis, coupled with folding analysis involving conserved allosteric hubs and regulatory phenylalanine residues, elucidates the identity of hydrophobic residues that have undergone minimal evolutionary changes over a span of 500 million years of apoptotic caspase evolution and which serve to stabilize this partially folded intermediate (Fig. 5A). In addition, MD simulations of unfolding in urea reveal that these residues collapse early in refolding, thus they exhibit characteristic features of a folding nucleus (Supplemental Video 1). Collectively, these findings demonstrate the presence of this intermediate, with a free energy of ∼2.7 kcal/mol, within the conformational ensemble of all apoptotic caspases and is depicted as the first high energy intermediate in the folding funnel (Fig. 6D).

Folding investigations with mutants of conserved allosteric hub residues, namely H304A and K351A + K353A (Fig. 6B) suggest that conserved allosteric hub residues shift the conformational equilibrium toward partially folded conformations in response to post translational modifications or other allosteric modulators in the allosteric pocket (Fig. 1B). Our folding studies involving mutations of highly conserved phenylalanine residues (caspase-8 F310A, F340A, F355A, and F399A) within the hydrophobic core, also result in a shift of the ensemble towards a partially folded intermediate that exhibits a conformational free energy of ∼2.4 kcal mol^-1^ (Fig. 6A and 6B), which may represent the same partially folded intermediate (∼2.7 kcals mol^-1^) observed in other mutants in this study (Supplemental Table S1). This similarity in conformational stabilities of the partially folded intermediate, observed in the various mutant sets, further adds evidence to the conclusion that high DC and BC residues (Fig. 5A) form the folding nucleus. Furthermore, the residues in the folding nucleus are also seen to be dynamic as the protein conformations vary (Supplemental Video 2), which we hypothesize to be related to the regulation of side chain packing within the hydrophobic core. Intriguingly, our findings suggest that, rather than playing a purely stabilizing role, these highly conserved phenylalanine residues, along with other conserved cysteine residues in the core, serve as a relay system for communication by facilitating the rapid transfer of charges from the allosteric pocket to the core or to the active site [40]. These insights not only make caspases an intriguing paradigm to further investigate electron tunnels in proteins, but they also provide a new avenue for investigating the mechanisms of allosteric communication in protein families and designing the next generation of small compounds for precise control of caspases.

## Materials and Methods

### Ancestral protein reconstruction (APR)

To resurrect a highly probable sequence of the last common ancestral caspase of the chordates involved the extrinsic pathway of apoptosis, we utilized a database of curated caspase sequences from CaspBase [42] that provided sequences from the initiator (caspase-8/-10/cFlip) and the effector (caspase-3/-6/-7) subfamilies in the chordate lineage. A total of 600 sequences were obtained to generate three databases comprising 200 sequences each for APR. Representative taxa from various classes of chordata (mammals, birds, fish, amphibians and reptiles) were chosen in each database to resurrect three probable ancestral sequences (AOA1, AOA2, and AOA3). Since the pro-domain is subject to high sequence variation due to recombinations, insertions, and deletions, we pruned the sequences on Jalview [43] to remove the pro-domains after PROMALS3D structure-based alignment in PROMALS3D [44]. Ancestral protein reconstruction was carried out as previously described [22].

### Homology modelling

For MD simulations of caspases in the enzymatically active configuration, we used mature caspases from the protein data bank (PDB): caspase-3 (PDB ID 3DEI), caspase-7 (PDB ID 1K86), caspase-6 (PDB ID 3NKF), caspase-8 (PDB ID 3KJQ), AOE-1 (PDB ID 6PDQ), and pseudo enzyme cFLIP (PDB ID 3H11) [45–49]. The rest of the ancestral enzymes were modelled based on the nearest available mature caspase structure in the evolutionary tree; for AOA-1,2,3 and for AOI-1. The ancestor of caspase-3/7 and of caspase-6 were modeled based on the crystal structures of caspase-3 (PDB ID 3dei) and of caspase-6 (PDB ID 3NKF), respectively. The ancestor of caspase-8/10 and caspase-10 were modeled based on the crystal structure of caspase-8 (PDB ID 3KJQ), and the ancestor of cFLIP was modeled based on the crystal structure of cFLIP(PDB ID 3H11). Procaspase enzymes from the PDB, exhibiting inactive but dimeric structures, were utilized for the dimeric conformation: procaspase-3 (PDB ID 4JQY), procaspase-7(PDB ID 1K88), procaspase-6(4N5D), and procaspase-8 (PDB ID 6px9) [46, 49–52]. Ancestor of caspase 3/7, AOE-1,2, and of AOA-1,2,3 were modelled using procaspase-3 (PDB ID 4JQY) as template, while ancestor of caspase-6 was modelled using procaspase-6 (PDB ID 4N5D) as template. We used the procaspase-8 (PDB ID 6PX9) as a template for the entire initiator tree from AOI-1,2 to the extant caspases. For the monomeric configuration, we modeled the complete tree using the procaspase-8 structure (PDB ID 2K7Z), the only NMR solution structure available [23].

### Network analysis of amino acid interactions

To analyze the ancestral caspase networks, we utilized the open-source software Cytoscape [53]. Through SenseNet [20],a Cytoscape plugin, we visualized and allocated measures of importance to amino acids by converting MD interaction timelines into protein structure networks. The networks were analyzed for degree and betweenness centrality for MD simulations performed in water and in 8 M urea. Simulations were imported onto SenseNet, and input parameters were modified to examine Van der Waals, hydrophobic, and electrostatic interactions with a distance cut-off of 4Å. The interaction weights were set to sum and the average, in order to generate network interactions displaying the average degree centrality (DC) and betweenness centrality (BC) values for each residue derived from the entire simulation. Nodes displaying high DC and BC were further classified according to the conservation scores obtained from Consurf [29].

### eMap electron pathway analysis

All protein structures obtained from the PDB and used in this study, including caspase-3 (PDB ID: 3DEI), caspase-7 (PDB ID: 1K86), caspase-6 (PDB ID: 3NKF), caspase-8 (PDB ID: 3KJQ), AOE-1 (PDB ID: 6PDQ), pseudo-enzyme cFLIP (PDB ID: 3H11), procaspase-6 (PDB ID: 4N5D), procaspase-8 (PDB ID: 6PX9), and the NMR solution structure of procaspase-8 (PDB ID: 2K7Z), were subjected to eMap analysis (https://emap.bu.edu/multiple) to identify shared electron and hole hopping pathways [39]. The proteins were used as inputs in the protein graph mining algorithm on the eMap web server to identify shared electron/hole hopping pathways among all the structures used in the study. Aromatic residues were selected for their electron transfer properties, along with cysteine and methionine due to their roles as electron relay centers [54]. Conserved pathways were identified and mapped onto the structures of caspase-8 (PDB ID: 6PX9 and PDB ID: 2K7Z), revealing common electron transfer networks.

### MD simulations and free energy landscape

MD simulations for 200 ns were performed in water and in 8M urea for all caspases shown in Supplemental Fig S1. Each caspase was examined in three conformations: the active, the dimeric, and the monomeric conformations, as previously described[15]. To study the concerted motions of caspases and to identify the most significant motions in the simulations, principal component analysis (PCA) was conducted for all protein atoms in the trajectory. The principal components (PCs) obtained from MD simulations in water and in urea are essentially the eigenvector values from the covariance matrix, each corresponding to a change in the trajectory. The eigenvalues and eigenvectors were analyzed using the gmx anaeig tool and the principal components (PCs) with the largest motions were selected and plotted for comparison. These PCs provide the main information about the spread of datapoints in the conformational space, indicating the global motion of the protein during simulations. To investigate the free energy landscape (FEL), the gmx sham tool was employed to combine the reaction coordinates of the PCs with the most significant movements. The FEL plots were generated using Matlab.

### Cloning, protein expression, and protein purification

The cloning, expression and purification for WT caspase-3 was carried out as previously described [13, 55]. Procaspase-3 and procaspase-8 network mutants (23 mutations) and other caspase-8 point mutants (caspase-8 H304A, caspase-8 K351A + K353A and caspase-8 F310A + F340A + F355A + F399A) were all cloned into pET21b+ plasmid, and sequences were confirmed by Sanger sequencing. Proteins were expressed in *E. coli* Lemo21(DE3) cells from NEB. We used constructs of caspase-8 where the pro-domain was removed to compare results with our previous studies [15]. Protein expression was carried out as previously described for procaspase-3, with the exception that we induced expression using 0.5 mM IPTG as the final concentration at 20 °C based on optimizing expression studies [55]. After 16 hours, cells were harvested by centrifugation at 5,000 rpm for 15 minutes, washed, and resuspended in buffer A (100 mM Tris-HCl,100 mM NaCl, pH 7.5). The cells were lysed using French press followed by centrifugation of lysate at 15,000 rpm to separate supernatant from cell debris. All the mutants appeared in the pellet, hence we performed denaturing purification by homogenizing the pellet in buffer B (6 M urea, 100 mM Tris-HCl,100 mM NaCl, pH 7.5) overnight and then centrifuging at 15,000 rpm to release the proteins from inclusion bodies. The supernatant was then incubated on His-bind resin for one hour, washed by 10 column volumes of buffer B, followed by gradient elution of imidazole from 0 mM to 500 mM in Buffer B. The samples were analyzed by SDS page and pure samples were collected. The samples were dialyzed and refolded overnight at 4 °C in buffer A containing 1 mM DTT and was then subjected to ion exchange chromatography (DEAE-sepharose) to separate impurities and further purify samples as previously described, with the exception that we used NaCl gradient instead of KCl gradient [55]. The samples were again dialyzed overnight at 4 °C in buffer A containing 1 mM DTT, concentrated, and stored at −80 °C.

### Folding studies preparation and data collection

Proteins were thawed on ice and subsequent folding and unfolding experiments were conducted in accordance with previously established protocols [56]. For unfolding and refolding studies, protein samples were prepared in phosphate buffer (pH 7.5, 1 mM DTT) with urea concentrations ranging from 0 to 9 M. Sample preparations and data collection using fluorescence emission spectroscopy and circular dichroism spectropolarimetry for unfolding and refolding studies were carried out as described before [15].

### Analysis and global fitting of data for equilibrium folding/unfolding

Unfolding and refolding data from fluorescence emission and circular dichroism studies were fit globally to the appropriate model as described previously [15, 56]. Briefly, we had a total of 10 datasets comprising folding data at 2 μM, 6 μM and 8 μM protein concentrations for each of the mutants. As there was no protein concentration dependence, the data were best fit to a monomeric three-state equilibrium folding model for caspase-3, caspase-8, and the caspase network mutants (23 mutants), as the data show a change in slope at ∼4 M urea (Fig 4B). In the three-state monomeric model, the native conformation (N) unfolds to a partially folded intermediate conformation (I) before fully unfolding (U).

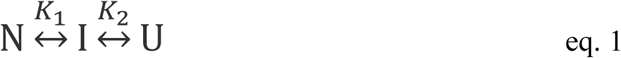

The data for the allosteric pocket mutants displayed a single transition and were best fit to a two-state folding model (Fig 6).

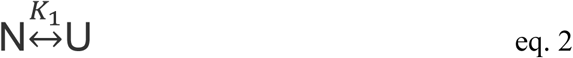

Detailed descriptions of both folding models have been previously documented [56]. Global fitting of data to the models described in the equations above was done utilizing Igor Pro (WaveMetrics, Inc). The global fitting is depicted as solid lines in Figure 4B and Figure 6A. The thermodynamic parameters, ΔG^°^ and m-values, from the global fits are presented in Supplemental Table S1.

## Supporting information

Supplemental Sequences

Video 1 Back

Video 1 Front

Video 2 Back

Video 2 Front

Video 3 Back

Video 3 Front

## Supplementary material description

Figure S1. Comparison of extant and ancestral caspases.

Figure S2. Violin plots representing the average degrees/contacts in caspases.

Figure S3. The free energy landscapes (FEL) of the dimeric conformation of caspases in water obtained from 200 ns MD simulations.

Figure S4. Target metastable states extracted from the last observed minima in the FEL of the dimeric conformation in 8 M urea.

Figure S5. The free energy landscapes of the monomeric conformation of caspases in urea obtained from 200 ns MD simulations.

Figure S6. MD average structures of the monomeric conformation in water.

Figure S7. Fluorescence emission and circular dichroism spectra of caspase-3 21M and of caspase-8 21M at pH 7, 25°C.

Figure S8. Limited trypsin proteolysis of caspase-3 WT, caspase-3 21M, and of caspase-8 21M.

Figure S9. Pair plots displaying the average degree centrality and average betweenness centrality.

Figure S10. Fluorescence emission and circular dichroism spectra of caspase-8 H304A, caspase-8 K351A + K353A, and of caspase-8 F310A + F340A + F355A + F399A at pH 7, 25°C.

Figure S11. Limited trypsin proteolysis of caspase-8 (H304A) (A) caspase-8 (K351A + K353A) (B), and of caspase-8 (F310A + F340A + F355A + F399A)

Figure S12. Results from eMap electron or hole transfer pathways analysis are mapped onto the monomeric and dimeric structures of caspase-8

Table S1. Conformational free energy for caspase-3 and caspase-8 mutants determined from folding/unfolding at pH 7.5.

Video 1 (Front and Back). The conformational transition of caspase-8 from a monomeric intermediate to a monomeric state, followed by a shift to a dimeric inactive conformation and finally to a dimeric active conformation. Key residues with high degree centrality (DC) and betweenness centrality (BC) are highlighted, with highly conserved residues shown in red and intermediately conserved residues in orange. The large subunit is shown in green and then small in cyan with the intersubunit linker in salmon. This video was generated by morphing average conformations of caspase-8 monomeric metastable state (Fig. 3C) in urea, procasaspe-8 monomeric conformation (PDB ID: 2k7z), procaspase-8 dimeric conformation (PDB ID: 6px9) and caspase-8 active dimeric conformation (PDB ID: 3kjq) in pymol.

Video 2 (Front and Back). Illustration of the dynamics near the allosteric hotspot and its interface with phenylalanine residues that link the allosteric hotspot to the network of amino acids throughout the rest of the caspase-8 structure during the transition from a monomeric to a dimeric conformation. Key residues highlighted: high DC and BC residues (red), phenylalanines F310, F340, F355, and F399 in the hydrophobic core (blue), the conserved N-terminal tyrosine Y226 (magenta), conserved proline P352 at the β-sheet 4 (light pink) linked to conserved lysines K351, K353 (yellow), and the 21 mutated residues in the small subunit (green). The morph video of procasaspe-8 monomeric conformation (PDB ID: 2k7z) and procaspase-8 dimeric conformation (PDB ID: 6px9) depicts a relay system for coordinating structural rearrangements, linking the allosteric hotspot to high DC and BC residues, active site (Histidine 317 and Cysteine C360 are shown as cyan side chains) and other dynamic networks via a cluster of phenyl alanine residues. Note that all residue numberings are in canonical capase-8 format.

Video 3 (Front and Back). Dynamics of high DC and BC residues driven by the allosteric hotspot and phenylalanine interface during the transition of caspase-8 from a monomeric to a dimeric conformation. High DC and BC residues are highlighted as spheres, while all other highlights and descriptions remain identical to those in video 2.

## Data availability

All data are contained in the article and supporting information.

## Author contributions

I.J., M.N.K.G., D.A.D. and A.C.C conceptualization and methodology; I.J., M.N.K.G., and D.A.D. investigation; I.J., M.N.K.G., D.A.D. and A.D. data fitting; I.J., M.N.K.G., D.A.D. and A.C.C writing-review and editing; A.C.C supervision.

## Funding and additional information

This work was supported by a grant from the National Institutes of Health (grant number: GM127654 [to A. C. C.]). The content is solely the responsibility of the authors and does not necessarily represent the official views of the National Institutes of Health.

