## Supplemental Figures for "Evolution of the conformational ensemble and allosteric networks of apoptotic caspases in chordates"

**Running title**: Evolutionary analysis of the conformational landscape of caspases

[*]Corresponding author: A. Clay Clark

[‡]Contributed equally

**Key Words**: caspase; network analysis; protein evolution; energy landscape, conformational dynamics, folding

### Supporting Information

**Supplemental Table S1.** Conformational free energy for caspase-3 and caspase-8 mutants determined from folding/unfolding at pH 7.5. The errors represent the standard deviation from the mean (mean±SD).

| **Enzyme** | **Folding Model** | **$\Delta G^0_1$ (kcal mol$^{-1}$)** | **$m_1$ (kcal mol$^{-1}$ M$^{-1}$)** | **$\Delta G^0_2$ (kcal mol$^{-1}$)** | **$m_2$ (kcal mol$^{-1}$ M$^{-1}$)** | **$\Delta G^0$ Total (kcal mol$^{-1}$)** | **m Total (kcal mol$^{-1}$ M$^{-1}$)** |
|---|---|---|---|---|---|---|---|
| Caspase-3 21M | N↔I↔U | 1.0 ± 0.01 | -0.1 ± 0.00 | 2.80 ± 0.30 | -0.31 ± 0.02 | 3.8 ± 0.01 | -0.41 ± 0.02 |
| Caspase-8 21M | N↔I↔U | 1.0 ± 0.03 | -0.2 ± 0.01 | 2.6 ± 0.41 | -0.48 ± 0.05 | 3.6 ± 0.02 | -0.7 ± 0.03 |
| Caspase-8 H304A | N↔U | | | 2.4 ± 0.23 | -0.76 ± 0.05 | 2.4 ± 0.23 | -0.76 ± 0.05 |
| Caspase-8 K351A K353A | N↔U | | | 2.4 ± 0.21 | -0.71 ± 0.05 | 2.4 ± 0.21 | -0.71 ± 0.05 |
| Caspase-8 F310A F340A F355A F399A | N↔U | | | 2.3 ± 0.24 | -0.90 ± 0.07 | 2.3 ± 0.24 | -0.90 ± 0.07 |

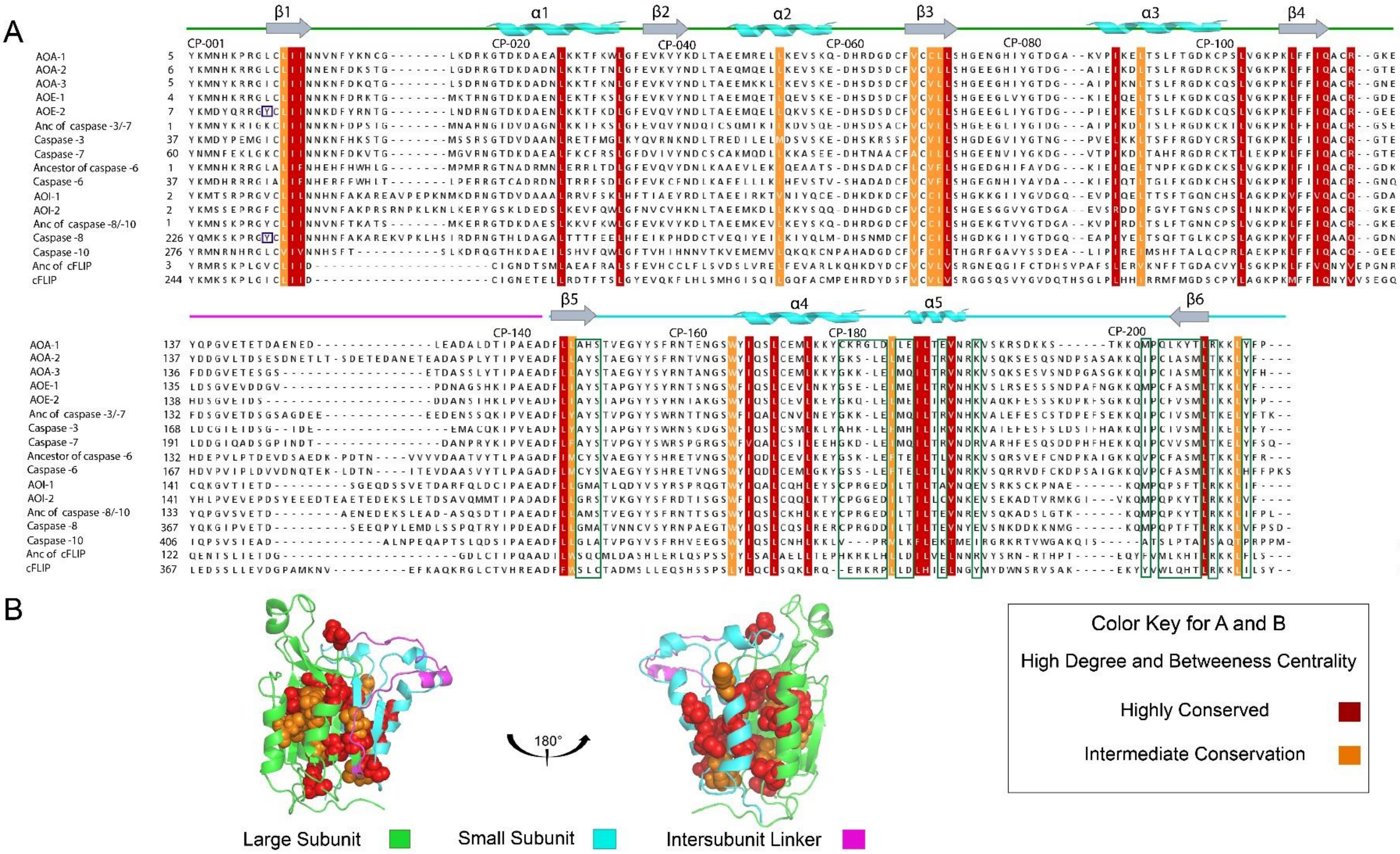


**Supplemental Figure S1.** Comparison of extant and ancestral caspases. (A) Multiple sequence alignment showing the secondary structural elements (loops, β-sheets, α-helices) along with the common position numbers among the caspases. The colored residues represent the amino acids that have high DC, BC and are classified as conserved residues (red) or intermediately conserved residues (orange) based on the conservation scores across the entire family. The residues that have different biochemical properties are boxed in green; the identified tyrosine residues close to H304 (caspase-8 numbering) is boxed in blue for caspase-8 and for AOE-2. The color of lines above the alignment correspond to the color scheme at the bottom. (B) High DC, BC residues classified as highly conserved (red spheres) or intermediately conserved (orange spheres) are mapped onto the dimeric conformation of caspase-8.

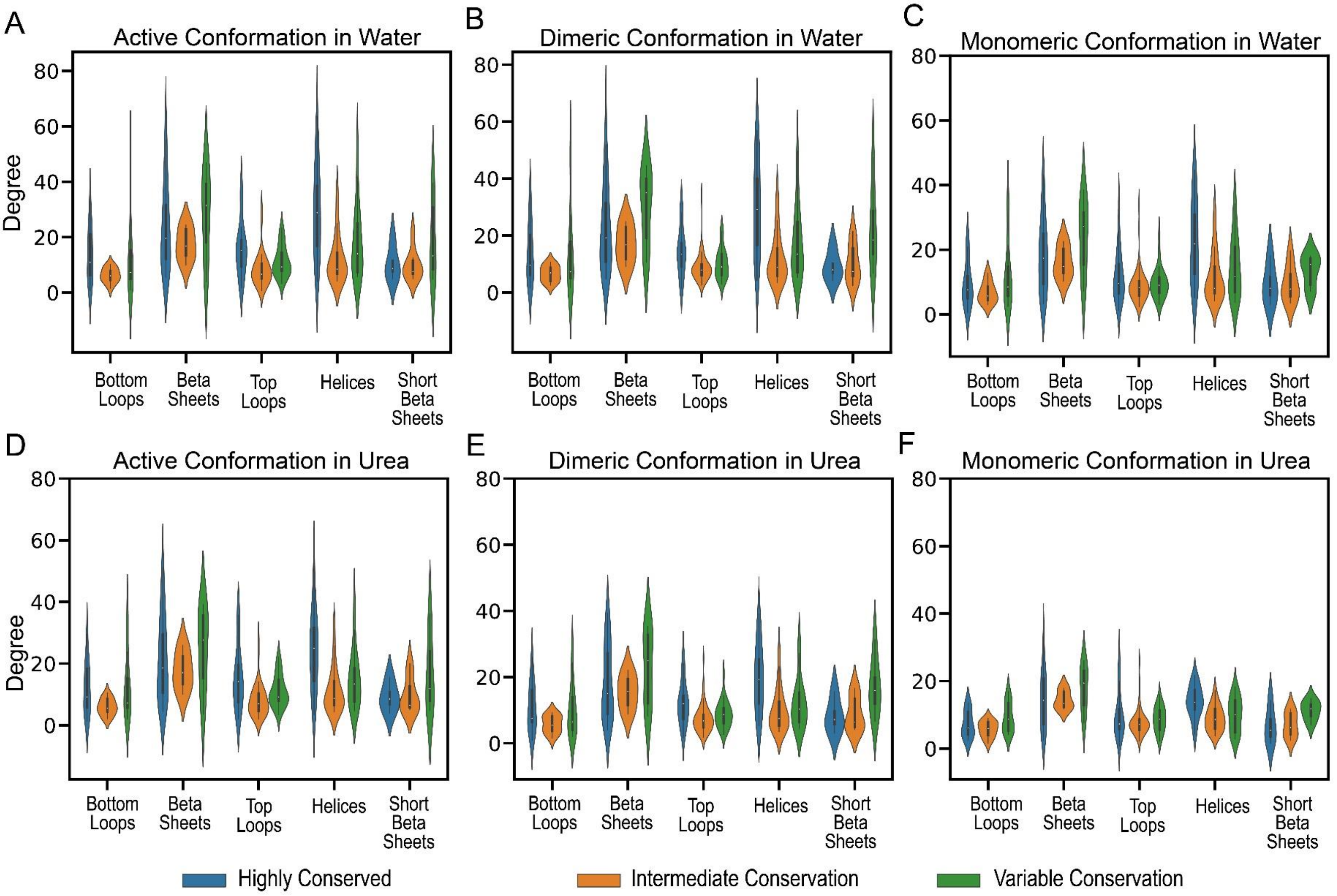


**Supplemental Figure S2.** Violin plots representing the average degrees/contacts categorized as conserved (blue), intermediately conserved (green), and variable (orange) based on the conservation scores and grouped according to secondary structural elements for the (A) active, (B) dimeric and (C) monomeric conformations in water and (D) active, (E) dimeric (F) monomeric conformations in 8 M urea.

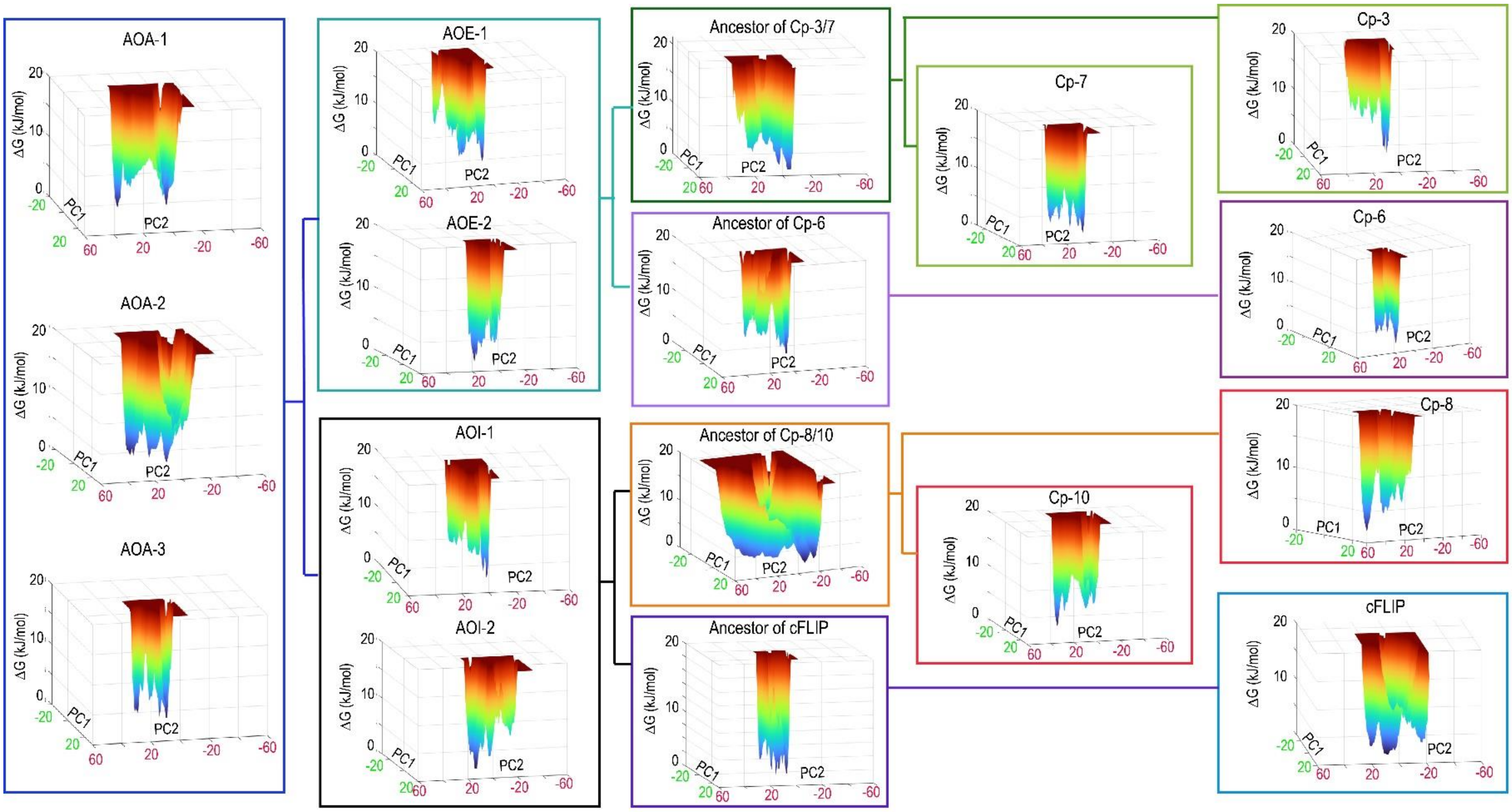


**Supplemental Figure S3.** The free energy landscapes (FEL) of the dimeric conformation of caspases in water obtained from 200 ns MD simulations. The FELs are generated as a function of projections of the MD trajectory onto the first (PC1) and the second (PC2) eigenvectors, respectively.

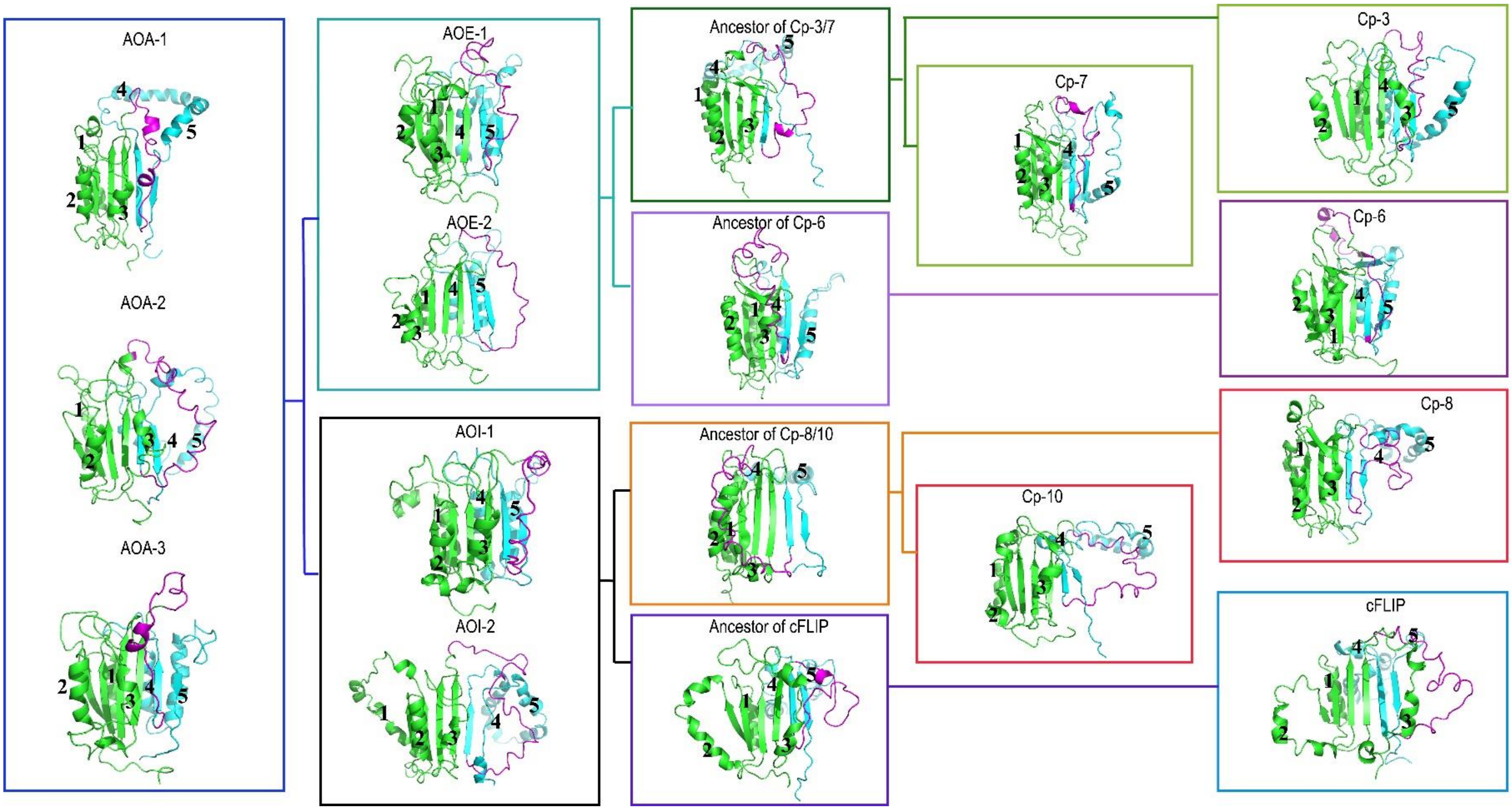


**Supplemental Figure S4.** Target metastable states extracted from the last observed minima in the free energy landscape (FEL) of the dimeric conformation in 8 M urea. The large subunit is highlighted in green, small subunit in cyan, and the inter-subunit linker in magenta.

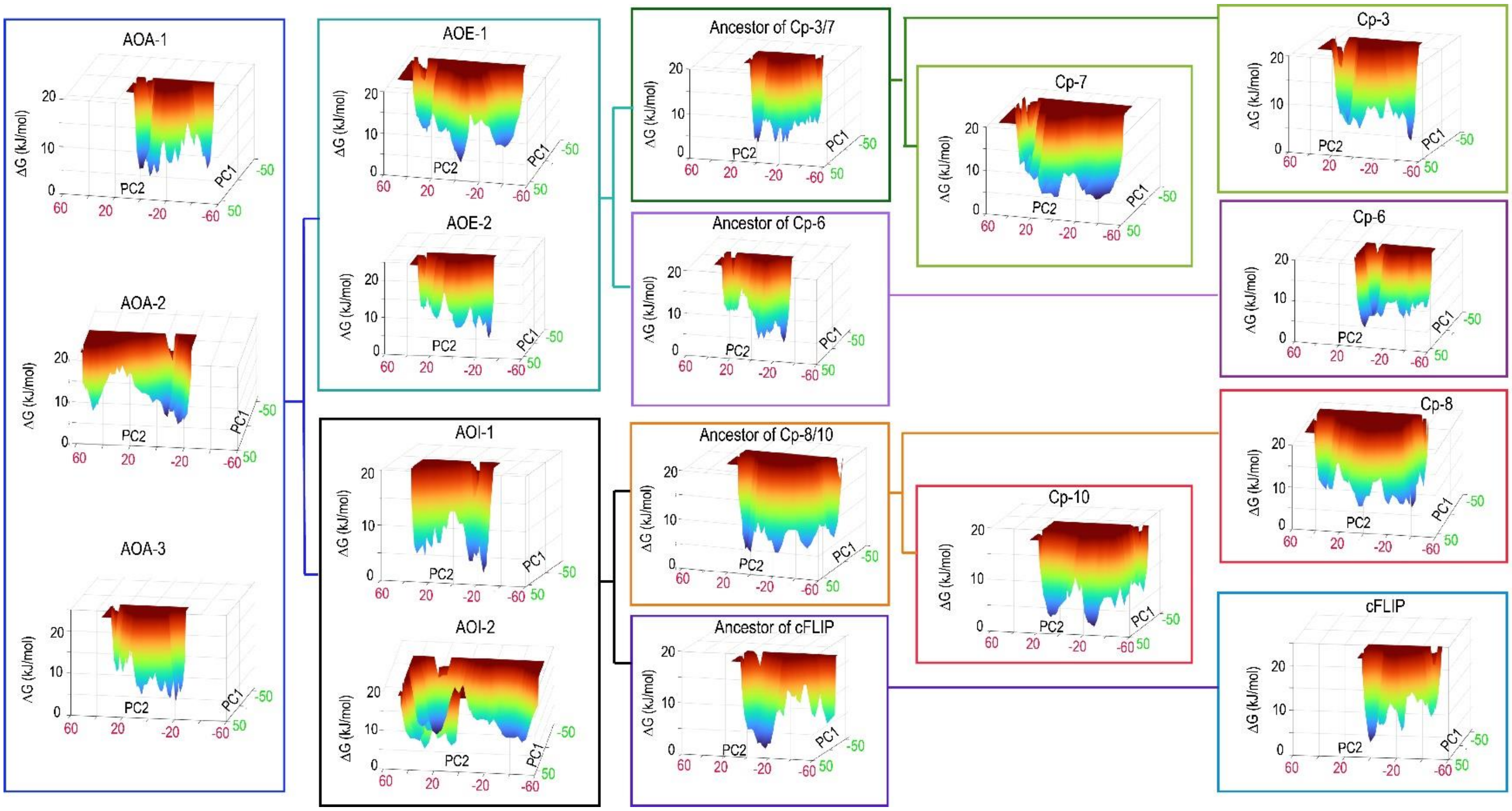


**Supplemental Figure S5.** The free energy landscapes (FEL) of the monomeric conformation of caspases in urea obtained from 200 ns MD simulations. The FELs are generated as a function of projections of the MD trajectory onto the first (PC1) and the second (PC2) eigenvectors, respectively.

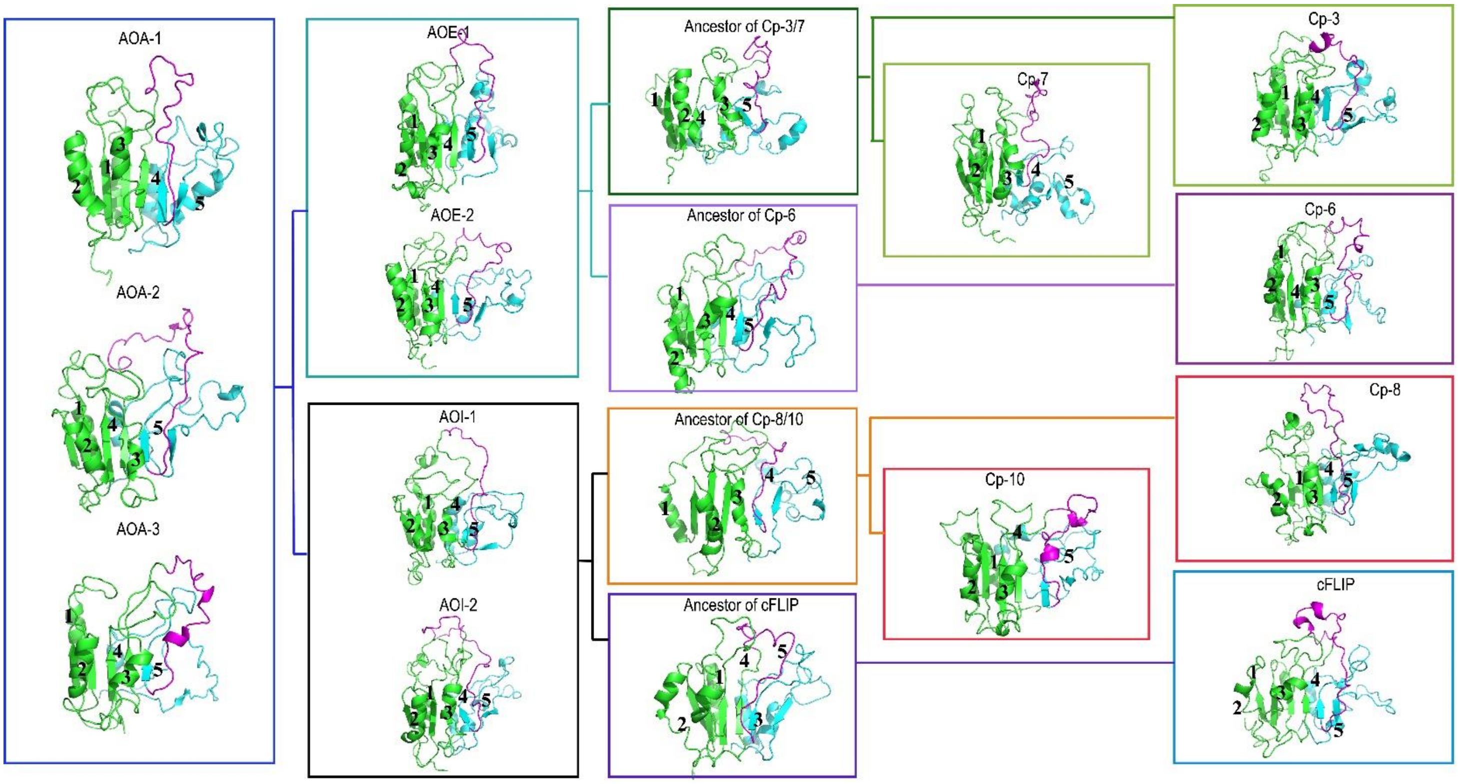


**Supplemental Figure S6.** Average structures obtained from MD simulations of the monomeric conformation in water. The large subunit is highlighted in green, small subunit in cyan, and the inter-subunit linker in magenta.

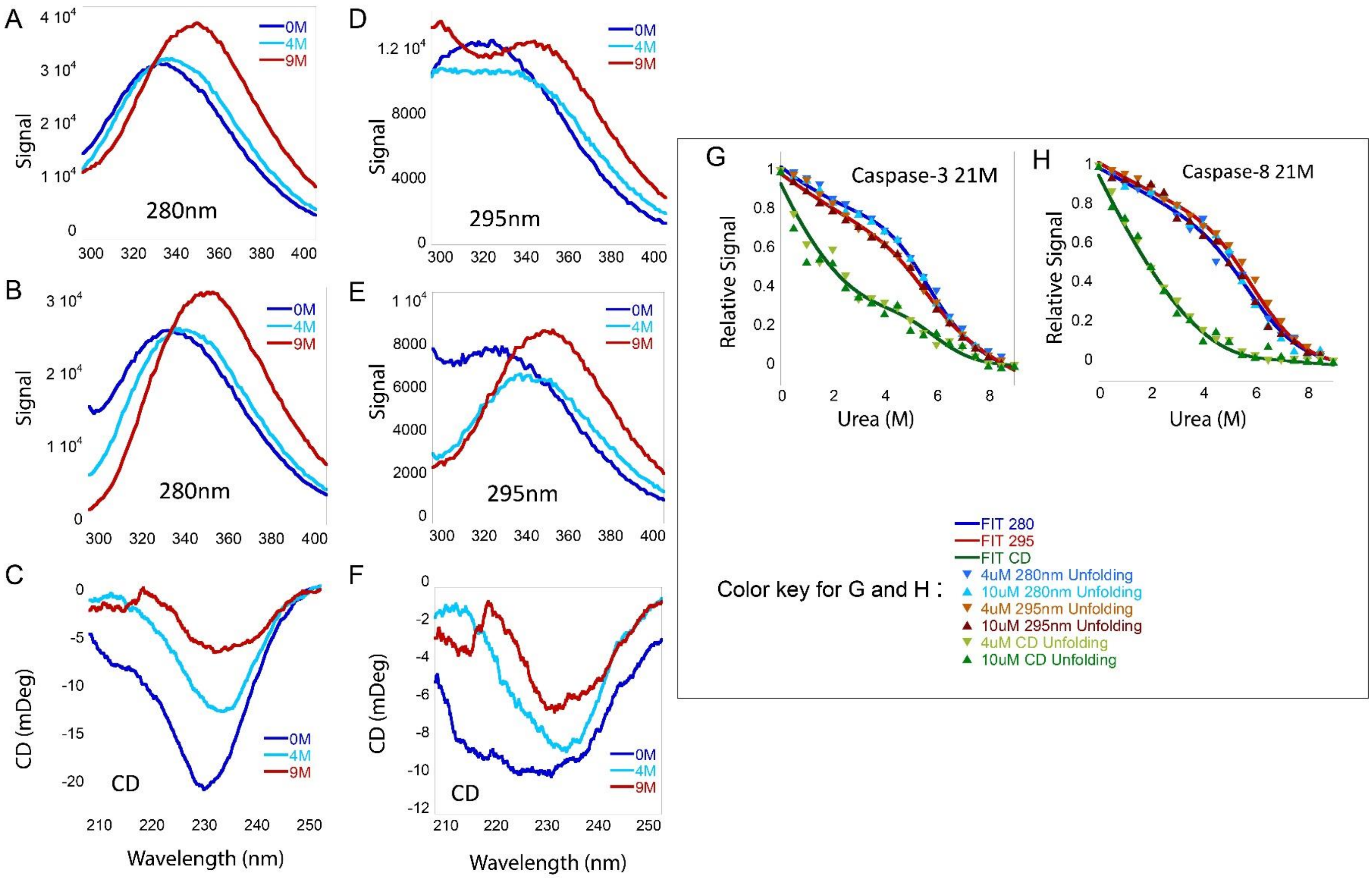


**Supplemental Figure S7**. Fluorescence emission and circular dichroism spectra of caspase-3 21M and of caspase-8 21M at pH 7, 25°C. Fluorescence emission spectra of caspase-8 21M following excitation at 280 nm (A) or 295 nm (D), and CD spectra (C). Fluorescence emission spectra of caspase-3 21M following excitation at 280 nm (B) or 295 nm (E), and CD spectra (F). Equilibrium unfolding of caspase-3 network mutant and caspase-8 network mutant at pH 7.5 monitored by fluorescence emission with excitation at 280 nm (●), 295 nm (●), CD (●) and refolding at 280 nm (▲), 295 nm (▲) and CD (▲), As described in the text, solid lines represent global fits to the data at 280 nm (▬), 295 nm (▬) and CD (▬), panels G and H, respectively.

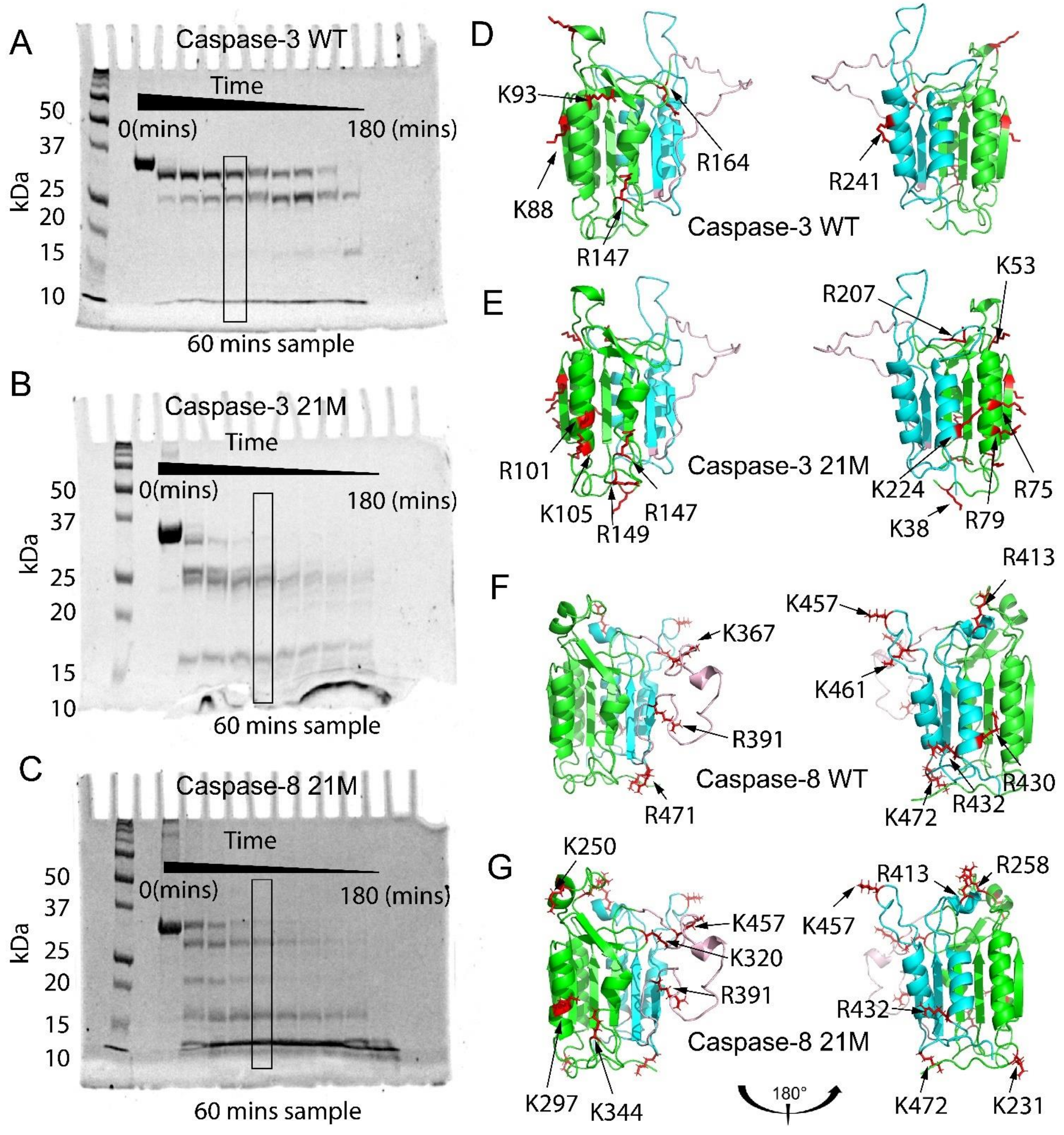


**Supplemental Figure S8**. Limited trypsin proteolysis of caspase-3 wild-type (A), caspase-3 21M (B), and of caspase-8 21M (C). Molecular weight markers are shown in the first lane on the gel, and the boxed seventh lane refers to the cleavage products that were analyzed. Cleavage products of limited trypsin proteolysis at pH 7.5 were determined by MALDI-TOF MS as described in the text and are represented on structures of caspase-3 wild-type (D) and 21M variant (E), and on caspase-8 wild-type (F) and 21M variant (G). The large subunit is depicted in green, the small subunit in cyan, inter-subunit linker in magenta, and the cleavage sites as red sticks.

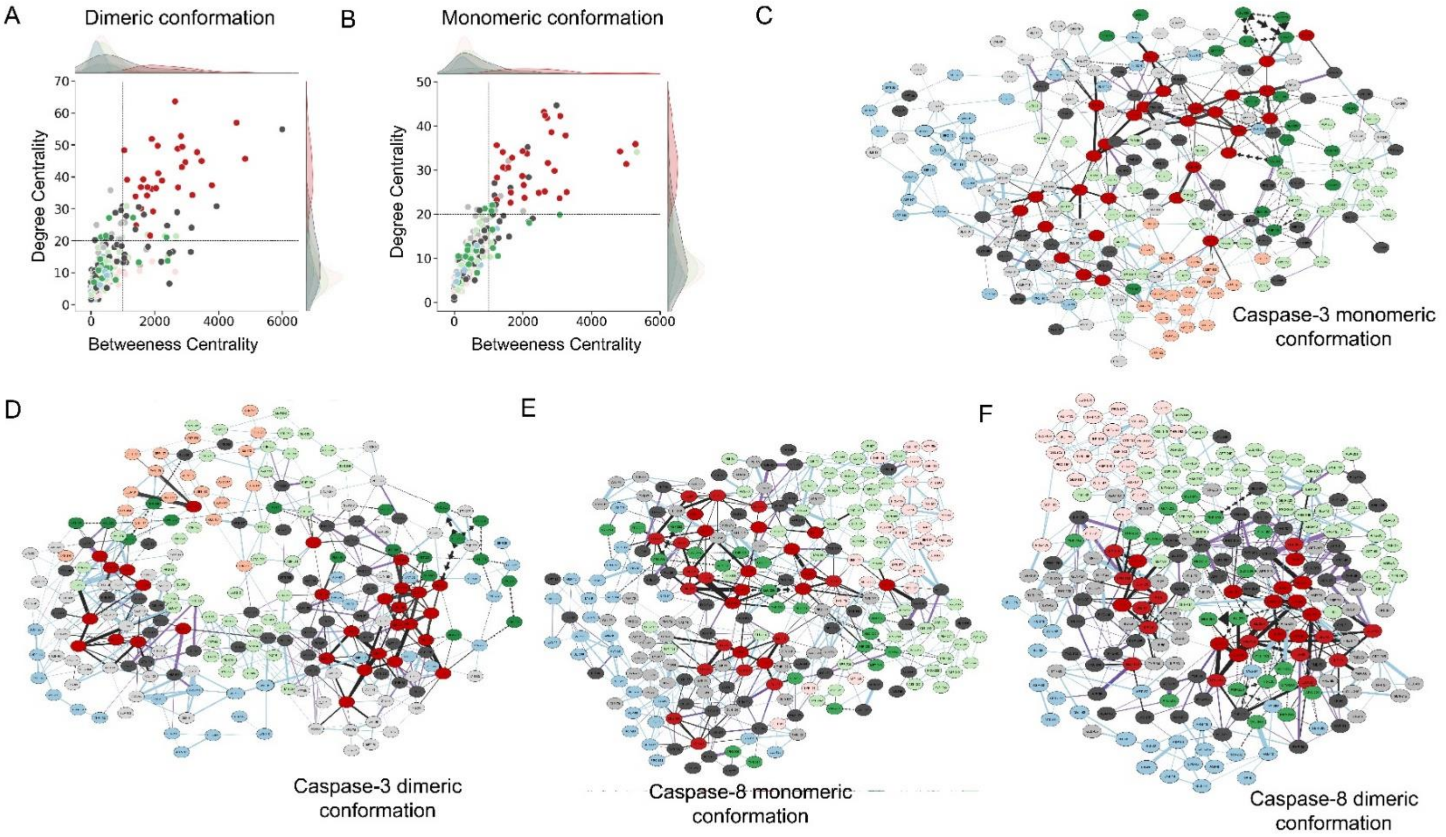


**Supplemental Figure S9**. Pair plots displaying the average degree centrality (DC) on y-axis and average betweenness centrality (BC) on x-axis for (A) dimeric conformation and (B) monomeric conformation in water. A 2D representation of the amino acid interaction networks for caspase-3 monomeric (C) and dimeric (D) conformations and for caspase-8 monomeric (E) and dimeric conformations (F) derived from MD simulations in water. The nodes on the network maps are color-coded according to the pair plots, with residues having a high degree and betweenness centrality colored red and show higher interactions (thick black lines) with the surrounding nodes. The 21M mutations are shown in green, highly conserved residues are shown in dark grey, the active site loops are shown in light green, the inter-subunit linker is shown in light pink, and the loops at the bottom of the structure are shown in light blue. All other elements are shown in light grey. Edges from high DC BC residues are shown as black edges; edges with source as any of the residues that are part of 21M mutants are shown as arrows.

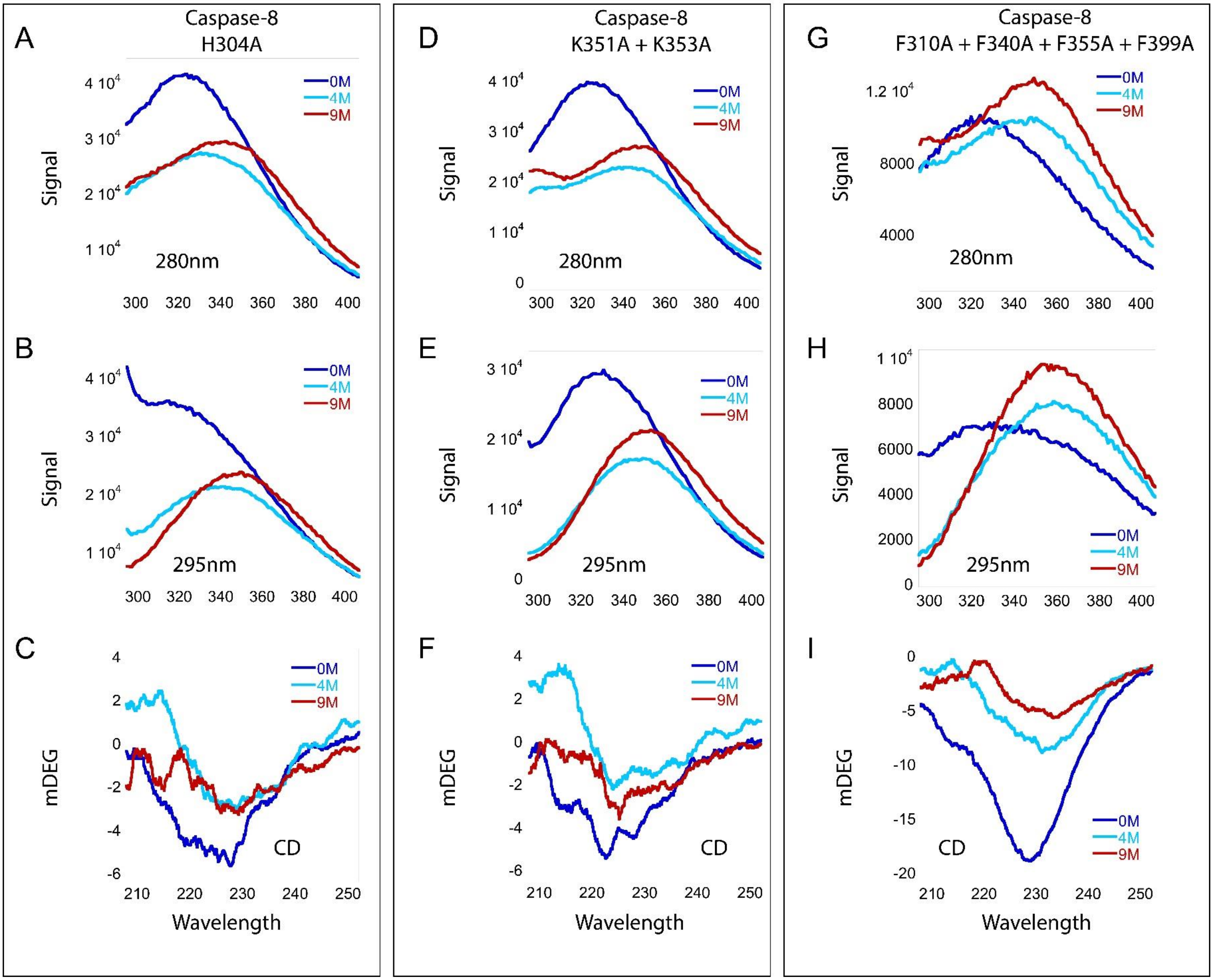


**Supplemental Figure S10.** Fluorescence emission and circular dichroism spectra of caspase-8 H304A, caspase-8 K351A + K353A, and caspase-8 F310A + F340A + F355A + F399A at pH 7, 25°C. Fluorescence emission spectra of caspase-8 H304A following excitation at 280 nm (A) or 295 nm (B), and CD spectra (C). Fluorescence emission spectra of caspase-8 H304A, following excitation at 280 nm (D) or 295 nm (E), and CD spectra (F). Fluorescence emission spectra of caspase-8 F310A + F340A + F355A + F399A, following excitation at 280 nm (G) or 295 nm (H), and CD spectra (I).

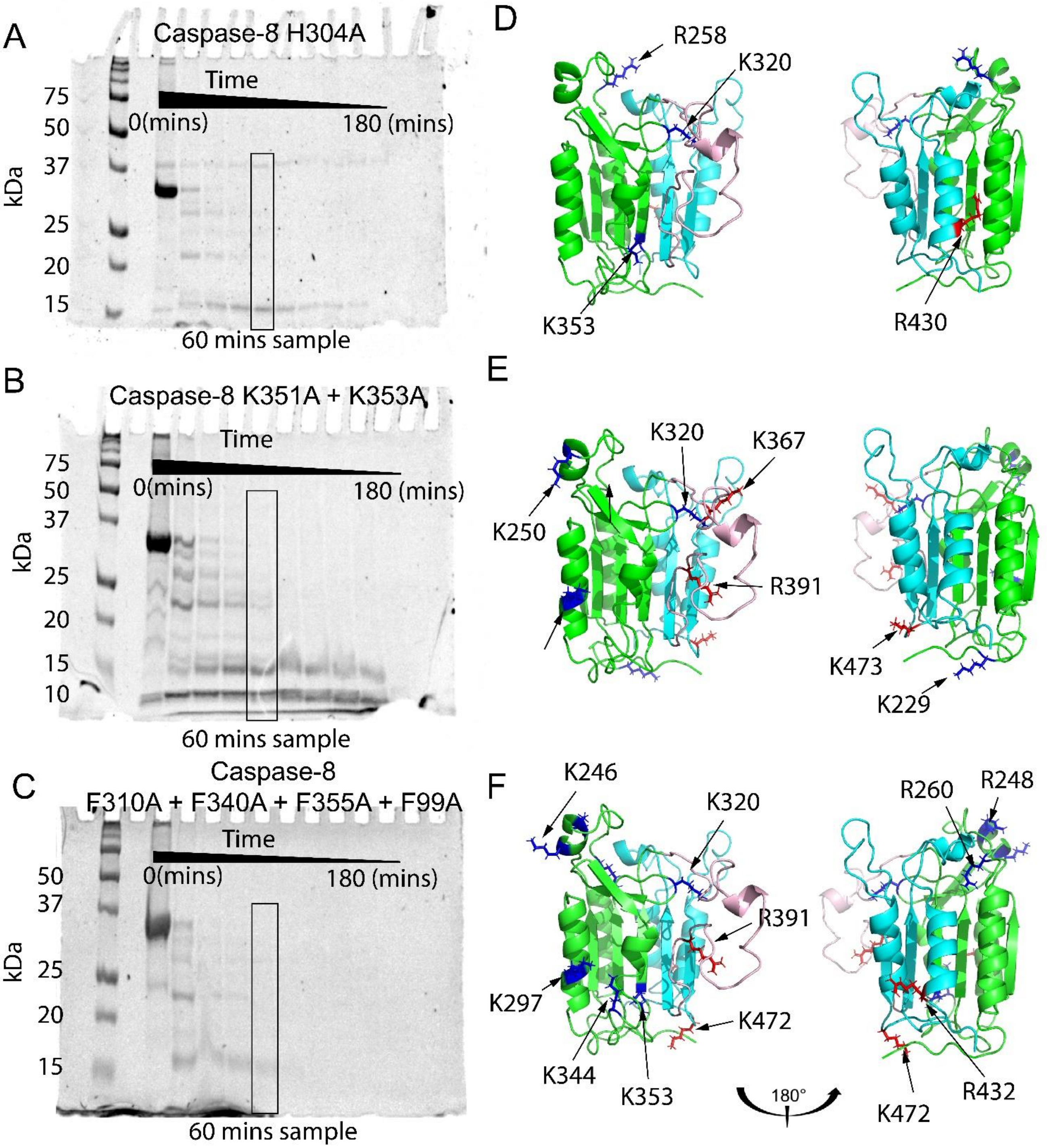


**Supplemental Figure S11.** Limited trypsin proteolysis of caspase-8 H304A (A) caspase-8 K351A + K353A (B), and of caspase-8 F310A + F340A + F355A + F399A (C). Molecular weight markers are shown in the first lane on the gel, and the boxed seventh lane refers to the cleavage products that were analyzed. Cleavage products of limited trypsin proteolysis at pH 7.5 were determined by MALDI-TOF MS as described in the text and represented on caspase-8 H304A (D), caspase-8 K351A + K353A (E) and caspase-8 F310A + F340A + F355A + F399A (F). The large subunit is depicted in green, the small subunit in cyan, inter-subunit linker in magenta, and the cleavage sites on the large subunit as blue and cleavage sites on the small subunit as red sticks.

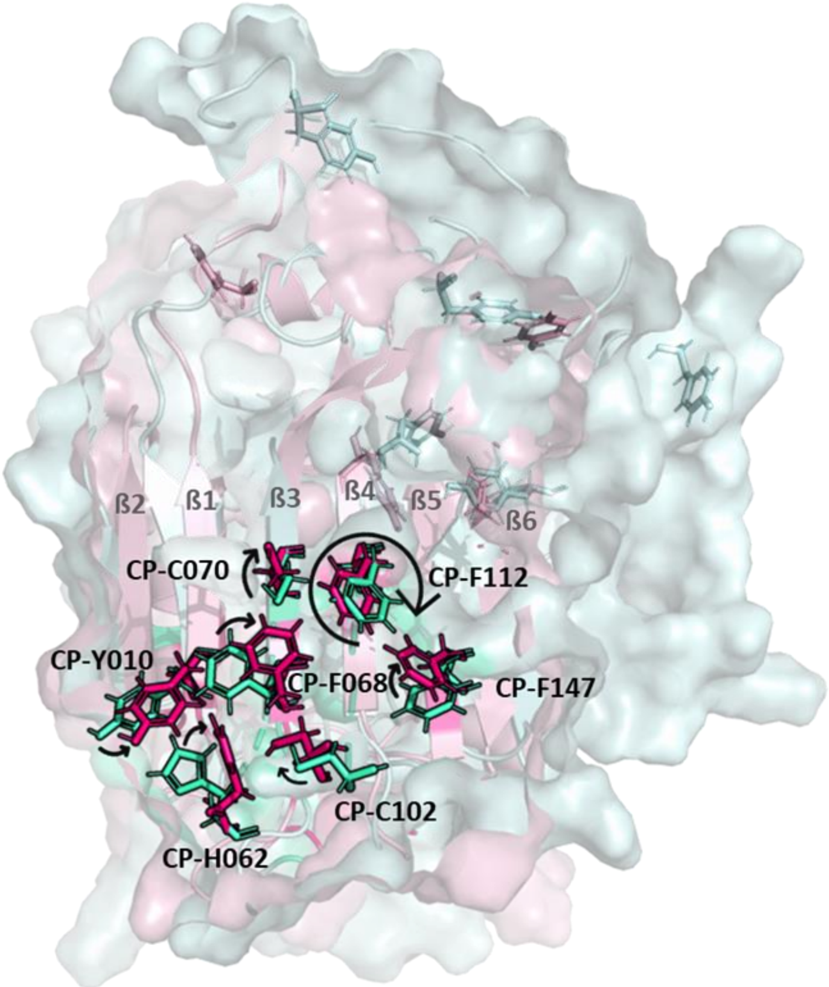


**Supplemental Figure S12.** Results from eMap electron or hole transfer pathways analysis are mapped onto the monomeric and dimeric structures of caspase-8 superimposed onto each other to show conformational dynamics of residues involved in electron or hole transport. The β-strands are labelled according to their order, whereas the residues are labeled according to the common position numbering (refer to Supplemental Fig S1)
