## Supplemental Sequences for "Evolution of the conformational ensemble and allosteric networks of apoptotic caspases in chordates"

**Running title**: Evolutionary analysis of the conformational landscape of caspases

[*]Corresponding author: A. Clay Clark

[‡]Contributed equally

**Key Words**: caspase; network analysis; protein evolution; energy landscape, conformational dynamics, folding

**Supporting Information**

| AOA1 | | | | |
|---|---|---|---|---|
| Accension ID | Caspase | Species | Class | Order |
| NP_004337.2 | Caspase-3 Isoform A Preprotein | Homo Sapiens | Mammalia | Primates |
| NP_001271338.1 | Caspase-3 | Mus Musculus | Mammalia | Rodentia |
| NP_571952.1 | Caspase-3 apoptosis-related cysteine peptidase a | Danio Rerio | Fish | Cypriniformes |
| XP_001517122.2 | Predicted: Caspase-3 | Ornithorhynchus Anatinus | Mammalia | Monotremata |
| NP_001003042.1 | Caspase-3 | Canis Lupus Familiaris | Mammalia | Carnivora |
| NP_001157433.1 | Caspase-3 | Equus Caballus | Mammalia | Perissodactyla |
| NP_990056.1 | Caspase-3 | Gallus Gallus | Birds | Galliformes |
| NP_999296.1 | Caspase-3 | Sus Scrofa | Mammalia | Artiodactyla |
| NP_001075586.1 | Caspase-3 | Oryctolagus Cuniculus | Mammalia | Lagomorpha |
| NP_001120900.1 | Caspase-3 | Xenopus Tropicalis | Amphibia | Anura |
| XP_011362327.1 | Predicted: Caspase-3 | Pteropus Vampyrus | Mammalia | Chiroptera |
| XP_003773079.1 | Predicted: Caspase-3 | Sarcophilus Harrisii | Mammalia | Dasyuromorphia |
| XP_004481806.1 | Predicted: Caspase-3 | Dasypus Novemcinctus | Mammalia | Cingulata |
| XP_006128558.1 | Predicted: Caspase-3 isoform X1 | Pelodiscus Sinensis | Reptilia | Testudines |
| XP_005149203.1 | Predicted: Caspase-3 | Melopsittacus Undulatus | Birds | Psittaciformes |
| XP_005282031.1 | Predicted: Caspase-3 isoform X1 | Chrysemys Picta Bellii | Reptilia | Testudines |
| XP_004389725.1 | Predicted: Caspase-3 | Trichechus Manatus Latirostris | Mammalia | Sirenia |
| XP_005045177.1 | Predicted: Caspase-3 | Ficedula Albicollis | Birds | Passeriformes |
| XP_019336883.1 | Predicted: Caspase-3 | Alligator Mississippiensis | Reptilia | Crocodilia |
| XP_007517715.1 | Predicted: Caspase-3 | Erinaceus Europaeus | Mammalia | Erinaceomorpha |
| NP_001098140.1 | Caspase-3 | Oryzias Latipes | Fish | Beloniformes |
| XP_005500235.1 | Caspase-3 | Columba Livia | Birds | Columbiformes |
| XP_022539261.1 | Caspase-3 | Astyanax Mexicanus | Fish | Characiformes |
| XP_009810205.1 | Predicted: Caspase-3 | Gavia Stellata | Birds | Gaviiformes |
| XP_009958677.1 | Predicted: Caspase-3 | Leptosomus Discolor | Birds | Leptosomiformes |
| XP_008569758.1 | Predicted: Caspase-3 | Galeopterus Variegatus | Mammalia | Dermoptera |
| XP_009473093.1 | Predicted: Caspase-3 | Nipponia Nippon | Birds | Pelecaniformes |
| NP_001290581.1 | Caspase-3 | Esox Lucius | Fish | Esociformes |
| XP_013910223.1 | Predicted: Caspase-3 | Thamnophis Sirtalis | Reptilia | Squamata |
| NP_001188010.1 | Caspase-3 | Ictalurus Punctatus | Fish | Siluriformes |
| NP_001081225.1 | Caspase-3 | Xenopus Laevis | Amphibia | Anura |
| XP_017503748.1 | Predicted: Caspase-3 | Manis Javanica | Mammalia | Pholidota |

| | | | | |
|---|---|---|---|---|
| NP_001269823.1 | Caspase-3 | Oreochromis Niloticus | Fish | Perciformes |
| XP_020863340.1 | Caspase-3 | Phascolarctos Cinereus | Mammalia | Diprotodontia |
| XP_020658385.1 | Caspase-3 | Pogona Vitticeps | Reptilia | Squamata |
| NP_001217.2 | Caspase-6 Isoform Alpha Precursor | Homo Sapiens | Mammalia | Primates |
| XP_016807502.1 | Predicted: Caspase-6 isoform X1 | Pan Troglodytes | Mammalia | Primates |
| NP_033941.3 | Caspase-6 Precursor | Mus Musculus | Mammalia | Rodentia |
| NP_113963.2 | Caspase-6 | Rattus Norvegicus | Mammalia | Rodentia |
| NP_001018333.1 | Caspase-6 | Danio Rerio | Fish | Cypriniformes |
| XP_545022.4 | Caspase-6 | Canis Lupus Familiaris | Mammalia | Carnivora |
| NP_990057.1 | Caspase-6 | Gallus Gallus | Birds | Galliformes |
| NP_001030496.1 | Caspase-6 | Bos Taurus | Mammalia | Artiodactyla |
| XP_008265713.1 | Predicted: Caspase-6 isoform X1 | Oryctolagus Cuniculus | Mammalia | Lagomorpha |
| NP_001011068.1 | Caspase-6 | Xenopus Tropicalis | Amphibia | Anura |
| XP_003221840.2 | Predicted: Caspase-6 | Anolis Carolinensis | Reptilia | Squamata |
| XP_006109127.2 | Predicted: Caspase-6 | Myotis Lucifugus | Mammalia | Chiroptera |
| XP_002197219.1 | Predicted: Caspase-6 | Taeniopygia Guttata | Birds | Passeriformes |
| XP_007907939.1 | Predicted: Caspase-6 | Callorhinchus Milii | Fish | Chimaeriformes |
| XP_003972493.2 | Predicted: Caspase-6 | Takifugu Rubripes | Fish | Tetraodontiformes |
| XP_007437451.1 | Predicted: Caspase-6 | Python Bivittatus | Reptilia | Squamata |
| XP_004476229.1 | Predicted: Caspase-6 isoform X1 | Dasypus Novemcinctus | Mammalia | Cingulata |
| XP_005999642.1 | Predicted: Caspase-6 | Latimeria Chalumnae | Fish | Coelacanthiformes |
| XP_005148683.1 | Predicted: Caspase-6 | Melopsittacus Undulatus | Birds | Psittaciformes |
| XP_014268841.1 | Caspase-6 isoform X2 | Maylandia Zebra | Fish | Perciformes |
| XP_005797675.2 | Predicted: Caspase-6 | Xiphophorus Maculatus | Fish | Cyprinodontiformes |
| XP_005287972.1 | Predicted: Caspase-6 | Chrysemys Picta Bellii | Reptilia | Testudines |
| XP_004380293.1 | Predicted: Caspase-6 | Trichechus Manatus Latirostris | Mammalia | Sirenia |
| XP_019355646.1 | Predicted: Caspase-6 isoform X1 | Alligator Mississippiensis | Reptilia | Crocodilia |
| XP_005517572.1 | Predicted: Caspase-6 | Pseudopodoces Humilis | Birds | Passeriformes |
| XP_006162304.2 | Predicted: Caspase-6 | Tupaia Chinensis | Mammalia | Scandentia |
| XP_005515276.1 | Predicted: Caspase-6 isoform X1 | Columba Livia | Birds | Columbiformes |
| XP_007054543.1 | Predicted: Caspase-6 | Chelonia Mydas | Reptilia | Testudines |
| XP_009663699.1 | Predicted: Caspase-6 isoform X1 | Struthio Camelus Australis | Birds | Struthioniformes |
| XP_008497831.1 | Predicted: Caspase-6 | Calypte Anna | Birds | Apodiformes |
| XP_011572791.1 | Predicted: Caspase-6 isoform X1 | Aquila Chrysaetos Canadensis | Birds | Accipitriformes |

| | | | | |
|---|---|---|---|---|
| XP_018425527.1 | Predicted: Caspase-6 | Nanorana Parkeri | Amphibia | Anura |
| XP_015281944.1 | Predicted: Caspase-6 isoform X1 | Gekko Japonicus | Reptilia | Squamata |
| NP_001081406.1 | Caspase-6 L Homeolog | Xenopus Laevis | Amphibia | Anura |
| NP_001117743.1 | Caspase-6 Precursor | Oncorhynchus Mykiss | Fish | Salmoniformes |
| NP_001253985.1 | Caspase-7 Isoform Alpha Precursor | Homo Sapiens | Mammalia | Primates |
| NP_071596.1 | Caspase-7 | Rattus Norvegicus | Mammalia | Rodentia |
| NP_001018443.1 | Caspase-7 | Danio Rerio | Fish | Cypriniformes |
| XP_001513388.4 | Predicted: Caspase-7 | Ornithorhynchus Anatinus | Mammalia | Monotremata |
| XP_005637795.1 | Caspase-7 isoform X1 | Canis Lupus Familiaris | Mammalia | Carnivora |
| XP_421764.3 | Predicted: Caspase-7 | Gallus Gallus | Birds | Galliformes |
| XP_020928981.1 | Caspase-7 isoform X3 | Sus Scrofa | Mammalia | Artiodactyla |
| XP_017204061.1 | Predicted: Caspase-7 isoform X2 | Oryctolagus Cuniculus | Mammalia | Lagomorpha |
| NP_001016299.1 | Caspase-7 | Xenopus Laevis | Amphibia | Anura |
| XP_008112945.1 | Predicted: Caspase-7 isoform X2 | Anolis Carolinensis | Reptilia | Squamata |
| XP_011357571.1 | Predicted: Caspase-7 isoform X1 | Pteropus Vampyrus | Mammalia | Chiroptera |
| XP_003961640.1 | Predicted: Caspase-7 | Takifugu Rubripes | Fish | Tetraodontiformes |
| XP_012396447.1 | Predicted: Caspase-7 | Sarcophilus Harrisii | Mammalia | Dasyuromorphia |
| XP_004458121.1 | Predicted: Caspase-7 | Dasypus Novemcinctus | Mammalia | Cingulata |
| XP_006002865.1 | Predicted: Caspase-7 isoform X1 | Latimeria Chalumnae | Fish | Coelacanthiformes |
| XP_006135034.1 | Predicted: Caspase-7 | Pelodiscus Sinensis | Reptilia | Testudines |
| XP_010331189.1 | Predicted: Caspase-7 | Saimiri Boliviensis Boliviensis | Mammalia | Primates |
| XP_004567546.1 | Caspase-7 isoform X3 | Maylandia Zebra | Fish | Perciformes |
| XP_005804467.1 | Predicted: Caspase-7 | Xiphophorus Maculatus | Fish | Cyprinodontiformes |
| XP_005293940.1 | Predicted: Caspase-7 | Chrysemys Picta Bellii | Reptilia | Testudines |
| XP_015203144.1 | Predicted: Caspase-7 isoform X1 | Lepisosteus Oculatus | Fish | Lepisosteiformes |
| XP_005048827.1 | Predicted: Caspase-7 isoform X2 | Ficedula Albicollis | Birds | Passeriformes |
| XP_014450146.1 | Predicted: Caspase-7 isoform X1 | Alligator Mississippiensis | Reptilia | Crocodilia |
| XP_004080311.2 | Caspase-7 | Oryzias Latipes | Fish | Beloniformes |
| XP_005520811.1 | Predicted: Caspase-7 | Pseudopodoces Humilis | Birds | Passeriformes |
| XP_021151155.1 | Caspase-7 isoform X1 | Columba Livia | Birds | Columbiformes |
| NP_001268771.1 | Caspase-7 | Mesocricetus Auratus | Mammalia | Rodentia |
| XP_005014501.1 | Caspase-7 | Anas Platyrhynchos | Birds | Anseriformes |
| XP_009965879.1 | Predicted: Caspase-7 | Tyto Alba | Birds | Strigiformes |

| | | | | |
|---|---|---|---|---|
| XP_010288794.1 | Predicted: Caspase-7 | Phaethon Lepturus | Birds | Phaethontiformes |
| XP_009483464.1 | Predicted: Caspase-7 | Pelecanus Crispus | Birds | Pelecaniformes |
| XP_018419981.1 | Predicted: Caspase-7 isoform X1 | Nanorana Parkeri | Amphibia | Anura |
| XP_018618189.1 | Predicted: Caspase-7 | Scleropages Formosus | Fish | Osteoglossiformes |
| NP_001081408.1 | Caspase-7 | Xenopus Laevis | Amphibia | Anura |
| NP_001091272.1 | Caspase-7 S Homeolog | Xenopus Laevis | Amphibia | Anura |
| NP_001219.2 | Caspase-8 Isoform A Precursor | Homo Sapiens | Mammalia | Primates |
| NP_001125222.2 | Caspase-8 | Pongo Abelii | Mammalia | Primates |
| NP_001264855.1 | Caspase-8 isoform 2 | Mus Musculus | Mammalia | Rodentia |
| NP_071613.1 | Caspase-8 | Rattus Norvegicus | Mammalia | Rodentia |
| XP_010600786.1 | Predicted: Caspase-8 isoform X1 | Loxodonta Africana | Mammalia | Proboscidea |
| NP_571585.2 | Caspase-8 | Danio Rerio | Fish | Cypriniformes |
| NP_001041494.1 | Caspase-8 | Canis Lupus Familiaris | Mammalia | Carnivora |
| XP_007501684.1 | Predicted: Caspase-8 isoform X1 | Monodelphis Domestica | Mammalia | Didelphimorphia |
| XP_014587812.1 | Predicted: Caspase-8 | Equus Caballus | Mammalia | Perissodactyla |
| NP_989923.1 | Caspase-8 | Gallus Gallus | Birds | Galliformes |
| NP_001026949.2 | Caspase-8 | Sus Scrofa | Mammalia | Artiodactyla |
| NP_001039435.1 | Caspase-8 | Bos Taurus | Mammalia | Artiodactyla |
| XP_017953067.1 | Predicted: Caspase-8 | Xenopus Tropicalis | Amphibia | Anura |
| XP_010711755.1 | Predicted: Caspase-8 | Meleagris Gallopavo | Birds | Galliformes |
| XP_014319897.1 | Predicted: Caspase-8 | Myotis Lucifugus | Mammalia | Chiroptera |
| XP_007882891.1 | Predicted: Caspase-8 | Callorhinchus Milii | Fish | Chimaeriformes |
| XP_011615013.1 | Predicted: Caspase-8 | Takifugu Rubripes | Fish | Tetraodontiformes |
| NP_001233725.1 | Caspase-8 | Cricetulus Griseus | Mammalia | Rodentia |
| XP_005306309.1 | Predicted: Caspase-8 isoform X1 | Chrysemys Picta Bellii | Reptilia | Testudines |
| XP_015214952.1 | Predicted: Caspase-8 | Lepisosteus Oculatus | Fish | Lepisosteiformes |
| XP_006272599.1 | Predicted: Caspase-8 isoform X1 | Alligator Mississippiensis | Reptilia | Crocodilia |
| XP_006272609.1 | Predicted: Caspase-8 | Alligator Mississippiensis | Reptilia | Crocodilia |
| NP_001098258.1 | Caspase-8 | Oryzias Latipes | Fish | Beloniformes |
| XP_012950190.1 | Predicted: Caspase-8 isoform X1 | Anas Platyrhynchos | Birds | Anseriformes |
| XP_009963925.1 | Predicted: Caspase-8 | Tyto Alba | Birds | Strigiformes |
| XP_009701337.1 | Predicted: Caspase-8 | Cariama Cristata | Birds | Cariamiformes |
| XP_009807896.1 | Predicted: Caspase-8 | Gavia Stellata | Birds | Gaviiformes |
| XP_017583733.1 | Predicted: Caspase-8 isoform X1 | Corvus Brachyrhynchos | Birds | Passeriformes |

| | | | | |
|---|---|---|---|---|
| XP_009938109.1 | Predicted: Caspase-8 | Opisthocomus Hoazin | Birds | Opisthocomiformes |
| XP_010017555.1 | Predicted: Caspase-8 | Nestor Notabilis | Birds | Psittaciformes |
| XP_009682548.1 | Predicted: Caspase-8 | Struthio Camelus Australis | Birds | Struthioniformes |
| XP_020795173.1 | Caspase-8 | Boleophthalmus Pectinirostris | Fish | Perciformes |
| XP_018585033.1 | Predicted: Caspase-8 isoform X1 | Scleropages Formosus | Fish | Osteoglossiformes |
| NP_001187127.1 | Caspase-8 | Ictalurus Punctatus | Fish | Siluriformes |
| NP_001079034.1 | Caspase-8 L Homeolog | Xenopus Laevis | Amphibia | Anura |
| NP_116759.2 | Caspase-10 Isoform 1 Preprotein | Homo Sapiens | Mammalia | Primates |
| NP_001127368.1 | Caspase-10 | Pongo Abelii | Mammalia | Primates |
| XP_007501670.1 | Predicted: Capase-10 | Monodelphis Domestica | Mammalia | Didelphimorphia |
| XP_005601738.1 | Predicted: Capase-10 | Equus Caballus | Mammalia | Perissodactyla |
| XP_421936.4 | Predicted: Capase-10 | Gallus Gallus | Birds | Galliformes |
| NP_001155112.1 | Caspase-10 | Sus Scrofa | Mammalia | Artiodactyla |
| NP_001093436.1 | Caspase-10 | Oryctolagus Cuniculus | Mammalia | Lagomorpha |
| NP_001015715.2 | Caspase-10 | Xenopus Tropicalis | Amphibia | Anura |
| XP_011218837.1 | Predicted: Caspase-10 isoform X1 | Ailuropoda Melanoleuca | Mammalia | Carnivora |
| XP_008118735.1 | Predicted: Caspase-10 isoform X1 | Anolis Carolinensis | Reptilia | Squamata |
| XP_010711745.1 | Predicted: Capase-10 | Meleagris Gallopavo | Birds | Galliformes |
| XP_006082464.1 | Predicted: Capase-10 | Myotis Lucifugus | Mammalia | Chiroptera |
| XP_011367468.1 | Predicted: Caspase-10 isoform X1 | Pteropus Vampyrus | Mammalia | Chiroptera |
| XP_003775343.2 | Predicted: Capase-10 | Sarcophilus Harrisii | Mammalia | Dasyuromorphia |
| XP_012917760.1 | Predicted: Capase-10 | Mustela Putorius Furo | Mammalia | Carnivora |
| XP_008175924.1 | Predicted: Caspase-10 isoform X3 | Chrysemys Picta Bellii | Reptilia | Testudines |
| XP_004378355.1 | Predicted: Capase-10 | Trichechus Manatus Latirostris | Mammalia | Sirenia |
| XP_005049234.1 | Predicted: Capase-10 | Ficedula Albicollis | Birds | Passeriformes |
| XP_004674884.1 | Predicted: Capase-10 | Condylura Cristata | Mammalia | Soricomorpha |
| XP_017802266.1 | Caspase-10 Isoform X1 | Papio Anubis | Mammalia | Primates |
| XP_014449789.1 | Predicted: Caspase-10 isoform X1 | Alligator Mississippiensis | Reptilia | Crocodilia |
| XP_014640222.1 | Predicted: Capase-10 | Ceratotherium simum simum | Mammalia | Perissodactyla |
| XP_012950189.1 | Caspase-10 | Anas Platyrhynchos | Birds | Anseriformes |
| XP_014374717.1 | Predicted: Caspase-10 isoform X1 | Alligator Sinensis | Reptilia | Crocodilia |
| XP_010288754.1 | Predicted: Capase-10 | Phaethon Lepturus | Birds | Phaethontiformes |

| | | | | |
|---|---|---|---|---|
| XP_009483191.1 | Predicted: Capase-10 | Pelecanus Crispus | Birds | Pelecaniformes |
| XP_009701338.1 | Predicted: Capase-10 | Cariama Cristata | Birds | Cariamiformes |
| XP_009924272.1 | Predicted: Capase-10 | Haliaeetus Albicilla | Birds | Accipitriformes |
| XP_009949374.1 | Predicted: Capase-10 | Leptosomus Discolor | Birds | Leptosomiformes |
| XP_008941675.1 | Predicted: Capase-10 | Merops Nubicus | Birds | Coraciiformes |
| XP_009938108.1 | Predicted: Capase-10 | Opisthocomus Hoazin | Birds | Opisthocomiformes |
| XP_010017544.1 | Predicted: Capase-10 | Nestor Notabilis | Birds | Psittaciformes |
| NP_001081410.1 | Caspase-10 S Homeolog | Xenopus Laevis | Amphibia | Anura |
| NP_001083130.1 | Caspase-10 L Homeolog | Xenopus Laevis | Amphibia | Anura |
| XP_019378751.1 | Predicted: Caspase-10 isoform X1 | Gavialis Gangeticus | Reptilia | Crocodilia |
| NP_001300701.1 | cFLIP | Danio Rerio | Fish | Cypriniformes |
| XP_008118737.1 | cFLIP | Anolis Carolinensis | Reptilia | Squamata |
| XP_034993049.1 | cFLIP | Zootoca Vivipara | Reptilia | Squamata |
| XP_015281011.1 | cFLIP | Gekko Japonicus | Reptilia | Squamata |
| XP_026709097.1 | cFLIP | Athene Cunicularia | Birds | Strigiformes |
| XP_009807246.1 | cFLIP | Gavia Stellata | Birds | Gaviiformes |
| XP_009479033.1 | cFLIP | Pelecanus Crispus | Birds | Pelecaniformes |
| XP_009282934.1 | cFLIP | Aptenodytes Forsteri | Birds | Sphenisciformes |
| XP_019378755.1 | cFLIP | Gavialis Gangeticus | Reptilia | Crocodilia |
| XP_006118011.1 | cFLIP | Pelodiscus Sinensis | Reptilia | Testudines |
| XP_007501666.1 | cFLIP | Monodelphis Domestica | Mammalia | Didelphimorphia |
| XP_031814208.1 | cFLIP | Sarcophilus harrisii | Mammalia | Dasyuromorphia |
| XP_019294730.1 | cFLIP | Panthera Pardus | Mammalia | Carnivora |
| XP_020930360.1 | cFLIP | Sus Scrofa | Mammalia | Artiodactyla |
| XP_032494111.1 | cFLIP | Phocoena Sinus | Mammalia | Cetacea |
| XP_027446588.1 | cFLIP | Zalophus Californianus | Mammalia | Carnivora |
| XP_019491241.1 | cFLIP | Hipposideros Armiger | Mammalia | Chiroptera |
| NP_001120655.1 | cFLIP | Homo Sapiens | Mammalia | Primates |
| NP_001125140.1 | cFLIP | Pongo Abelii | Mammalia | Primates |
| XP_028365650.1 | cFLIP | Phyllostomus Discolor | Mammalia | Chiroptera |
| NP_001188445.1 | cFLIP | Oryzias Latipes | Fish | Beloniformes |
| XP_022595948.1 | cFLIP | Seriola Dumerili | Fish | Perciformes |
| XP_026152299.1 | cFLIP | Mastacembelus Armatus | Fish | Synbranchiformes |
| XP_019945253.1 | cFLIP | Paralichthys Olivaceus | Fish | Pleuronectiformes |
| NP_001254595.1 | cFLIP | Gasterosteus Aculeatus | Fish | Gasterosteiformes |
| **AOA 2** | | | | |
| Accension ID | Caspase | Species | Class | Order |

| | | | | |
|---|---|---|---|---|
| NP_004337.2 | Caspase-3 isoform a preproprotein | Homo Sapiens | Mammalia | Primates |
| NP_001012435.1 | Caspase-3 isoform a preproprotein | Pan Troglodyte | Mammalia | Primates |
| NP_571952.1 | Caspase-3 apoptosis-related cysteine peptidase a | Danio Rerio | Fish | Cypriniformes |
| NP_001003042.1 | Caspase-3 | Canis Lupus Familiaris | Mammalia | Carnivora |
| NP_001157433.1 | Caspase-3 | Equus Caballus | Mammalia | Perissodactyla |
| NP_990056.1 | Caspase-3 | Gallus Gallus | Birds | Galliformes |
| NP_001071308.1 | Caspase-3 | Bos Taurus | Mammalia | Artiodactyla |
| NP_001075586.1 | Caspase-3 | Oryctolagus Cuniculus | Mammalia | Lagomorpha |
| NP_001120900.1 | Caspase-3 | Xenopus Tropicalis | Amphibia | Anura |
| XP_014314596.1 | Predicted: Caspase-3 isoform X1 | Myotis Lucifugus | Mammalia | Chiroptera |
| XP_004610417.1 | Predicted: Caspase-3 | Sorex Araneus | Mammalia | Soricomorpha |
| XP_014351567.1 | Predicted: Caspase-3 | Latimeria Chalumnae | Coelacanthi | Coelacanthiformes |
| XP_006128558.1 | Predicted: Caspase-3 isoform X1 | Pelodiscus Sinensis | Reptilia | Testudines |
| XP_004682433.1 | Predicted: Caspase-3 | Condylura Cristata | Mammalia | Soricomorpha |
| XP_004579072.1 | Predicted: Caspase-3 | Ochotona Princeps | Mammalia | Lagomorpha |
| XP_007939063.1 | Predicted: Caspase-3 | Orycteropus afer afer | Mammalia | Tubulidentata |
| XP_006882296.1 | Predicted: Caspase-3 | Elephantulus Edwardii | Mammalia | Macroscelidea |
| NP_001098168.1 | Caspase-3B | Oryzias Latipes | Fish | Beloniformes |
| XP_006916432.1 | Predicted: Caspase-3 | Pteropus Alecto | Mammalia | Chiroptera |
| XP_014114736.1 | Predicted: Caspase-3 | Pseudopodoces Humilis | Birds | Passeriformes |
| XP_005445415.1 | Predicted: Caspase-3 | Falco Cherrug | Birds | Falconiformes |
| XP_007054526.1 | Predicted: Caspase-3 | Chelonia Mydas | Reptilia | Testudines |
| XP_005030551.1 | Caspase-3 | Anas Platyrhynchos | Birds | Anseriformes |
| NP_001271764.1 | Caspase-3 | Macaca Fascicularis | Mammalia | Primates |
| XP_006026683.1 | Predicted: Caspase-3 | Alligator Sinensis | Reptilia | Crocodilia |
| XP_007549682.1 | Predicted: Caspase-3 isoform X1 | Poecilia Formosa | Fish | Cyprinodontiformes |
| XP_009970400.1 | Predicted: Caspase-3 | Tyto Alba | Birds | Strigiformes |
| XP_009479867.1 | Predicted: Caspase-3 | Pelecanus Crispus | Birds | Pelecaniformes |
| XP_009922022.1 | Predicted: Caspase-3 | Haliaeetus Albicilla | Birds | Accipitriformes |
| NP_001290581.1 | Caspase-3 | Esox Lucius | Fish | Esociformes |
| XP_015687286.1 | Predicted: Caspase-3 | Protobothrops Mucrosquamatus | Reptilia | Squamata |
| NP_001188010.1 | Caspase-3 | Ictalurus Punctatus | Fish | Siluriformes |
| NP_001081225.1 | Caspase-3 | Xenopus Laevis | Amphibia | Anura |
| NP_001273018.1 | Caspase-3 | Capra Hircus | Mammalia | Artiodactyla |
| NP_001269823.1 | Caspase-3 | Oreochromis Niloticus | Fish | Perciformes |

| | | | | |
|---|---|---|---|---|
| NP_001217.2 | Caspase-6 Isoform Alpha Precursor | Homo Sapiens | Mammalia | Primates |
| NP_033941.3 | Caspase-6 Precursor | Mus Musculus | Mammalia | Rodentia |
| NP_113963.2 | Caspase-6 | Rattus Norvegicus | Mammalia | Rodentia |
| XP_010588808.1 | Predicted: Caspase-6 | Loxodonta Africana | Mammalia | Proboscidea |
| NP_001018333.1 | Caspase-6 | Danio Rerio | Fish | Cypriniformes |
| NP_990057.1 | Caspase-6 | Gallus Gallus | Birds | Galliformes |
| NP_001030496.1 | Caspase-6 | Bos Taurus | Mammalia | Artiodactyla |
| NP_001011068.1 | Caspase-6 | Xenopus Tropicalis | Amphibia | Anura |
| XP_011359747.1 | Predicted: Caspase-6 Isoform X1 | Pteropus Vampyrus | Mammalia | Chiroptera |
| XP_015101393.1 | Predicted: Caspase-6 | Vicugna Pacos | Mammalia | Artiodactyla |
| XP_004748146.1 | Predicted: Caspase-6 Isoform X2 | Mustela Putorius Furo | Mammalia | Carnivora |
| XP_014427079.1 | Predicted: Caspase-6 | Pelodiscus Sinensis | Reptilia | Testudines |
| XP_006629925.2 | Predicted: Caspase-6 | Lepisosteus Oculatus | Holostei | Lepisosteiformes |
| XP_004380293.1 | Predicted: Caspase-6 | Trichechus Manatus Latirostris | Mammalia | Sirenia |
| XP_005044995.1 | Predicted: Caspase-6 Isoform X1 | Ficedula Albicollis | Birds | Passeriformes |
| XP_012583463.1 | Predicted: Caspase-6 Isoform X2 | Condylura Cristata | Mammalia | Soricomorpha |
| XP_006881156.1 | Predicted: Caspase-6 | Elephantulus Edwardii | Mammalia | Macroscelidea |
| XP_004084686.1 | Caspase-6 | Oryzias Latipes | Fish | Beloniformes |
| XP_012860317.1 | Predicted: Caspase-6 | Echinops Telfairi | Mammalia | Afrosoricida |
| XP_013155138.1 | Predicted: Caspase-6 | Falco Peregrinus | Birds | Falconiformes |
| XP_007054543.1 | Predicted: Caspase-6 | Chelonia Mydas | Reptilia | Testudines |
| XP_021128354.1 | Caspase-6 | Anas Platyrhynchos | Birds | Anseriformes |
| XP_006025610.2 | Predicted: Caspase-6 | Alligator Sinensis | Reptilia | Crocodilia |
| XP_007560684.1 | Predicted: Caspase-6 Isoform X1 | Poecilia Formosa | Fish | Cyprinodontiformes |
| XP_009966306.1 | Predicted: Caspase-6 | Tyto Alba | Birds | Strigiformes |
| XP_010290767.1 | Predicted: Caspase-6 | Phaethon Lepturus | Birds | Phaethontiformes |
| XP_009483599.1 | Predicted: Caspase-6 | Pelecanus Crispus | Birds | Pelecaniformes |
| XP_009699574.1 | Predicted: Caspase-6 | Cariama Cristata | Birds | Cariamiformes |
| XP_008285572.1 | Predicted: Caspase-6 | Stegastes Partitus | Fish | Perciformes |
| XP_019110884.1 | Predicted: Caspase-6 Isoform X1 | Larimichthys Crocea | Fish | Perciformes |
| XP_015666700.1 | Predicted: Caspase-6 | Protobothrops Mucrosquamatus | Reptilia | Squamata |
| XP_018611669.1 | Predicted: Caspase-6 | Scleropages Formosus | Fish | Osteoglossiformes |
| NP_001081406.1 | Caspase-6 L Homeolog | Xenopus Laevis | Amphibia | Anura |
| XP_019377505.1 | Predicted: Caspase-6 Isoform X1 | Gavialis Gangeticus | Reptilia | Crocodilia |

| | | | | |
|---|---|---|---|---|
| NP_001117743.1 | Caspase-6 Precursor | Oncorhynchus Mykiss | Fish | Salmoniformes |
| NP_001253985.1 | Caspase-7 isoform alpha precursor | Homo Sapiens | Mammalia | Primates |
| NP_071596.1 | Caspase-7 | Rattus Norvegicus | Mammalia | Rodentia |
| XP_010587277.1 | Predicted: Caspase-7 isoform X1 | Loxodonta Africana | Mammalia | Proboscidea |
| NP_001018443.1 | Caspase-7 | Danio Rerio | Fish | Cypriniformes |
| XP_421764.3 | Predicted: Caspase-7 | Gallus Gallus | Birds | Galliformes |
| XP_002698555.1 | Predicted: Caspase-7 | Bos Taurus | Mammalia | Artiodactyla |
| NP_001016299.1 | Caspase-7 | Xenopus Tropicalis | Amphibia | Anura |
| XP_011230112.1 | Predicted: Caspase-7 isoform X2 | Ailuropoda Melanoleuca | Mammalia | Carnivora |
| XP_008112945.1 | Predicted: Caspase-7 isoform X2 | Anolis Carolinensis | Reptilia | Squamata |
| XP_014310783.1 | Predicted: Caspase-7 | Myotis Lucifugus | Mammalia | Chiroptera |
| XP_004612023.2 | Predicted: Caspase-7 | Sorex Araneus | Mammalia | Soricomorpha |
| XP_006135034.1 | Predicted: Caspase-7 | Pelodiscus Sinensis | Reptilia | Testudines |
| XP_005154576.1 | Predicted: Caspase-7 isoform X1 | Melopsittacus Undulatus | Birds | Psittaciformes |
| XP_005747456.1 | Predicted: Caspase-7 isoform X1 | Pundamilia Nyererei | Actinopterygii | Cichliformes |
| XP_004681050.1 | Predicted: Caspase-7 | Condylura Cristata | Mammalia | Soricomorpha |
| XP_014450146.1 | Predicted: Caspase-7 isoform X1 | Alligator Mississippiensis | Reptilia | Crocodilia |
| XP_014641239.1 | Predicted: Caspase-7 | Ceratotherium simum simum | Mammalia | Perissodactyla |
| XP_006831548.1 | Predicted: Caspase-7 | Chrysochloris Asiatica | Mammalia | Afrosoricida |
| XP_005888715.1 | Predicted: Caspase-7 | Bos Taurus | Mammalia | Artiodactyla |
| XP_005520811.1 | Predicted: Caspase-7 | Pseudopodoces Humilis | Birds | Passeriformes |
| XP_005239289.1 | Predicted: Caspase-7 isoform X1 | Falco Peregrinus | Birds | Falconiformes |
| NP_001268771.1 | Caspase-7 | Mesocricetus Auratus | Mammalia | Rodentia |
| XP_005481147.1 | Predicted: Caspase-7 | Zonotrichia Albicollis | Birds | Passeriformes |
| XP_007574351.1 | Predicted: Caspase-7 isoform X1 | Poecilia Formosa | Fish | Cyprinodontiformes |
| XP_008312002.1 | Predicted: Caspase-7 isoform X1 | Cynoglossus Semilaevis | Fish | Pleuronectiformes |
| XP_009697487.1 | Predicted: Caspase-7 | Cariama Cristata | Birds | Cariamiformes |
| XP_010198191.1 | Predicted: Caspase-7 | Colius Striatus | Birds | Coliiformes |
| XP_008275786.1 | Predicted: Caspase-7 | Stegastes Partitus | Fish | Perciformes |
| XP_009578529.1 | Predicted: Caspase-7 | Fulmarus Glacialis | Birds | Procellariiformes |
| XP_009923386.1 | Predicted: Caspase-7 | Haliaeetus Albicilla | Birds | Accipitriformes |
| XP_010006222.1 | Predicted: Caspase-7 isoform X1 | Chaetura Pelagica | Birds | Apodiformes |

| | | | | |
|---|---|---|---|---|
| XP_018419981.1 | Predicted: Caspase-7 isoform X1 | Nanorana Parkeri | Amphibia | Anura |
| XP_012670088.1 | Predicted: Caspase-7 | Clupea Harengus | Fish | Clupeiformes |
| XP_018618189.1 | Predicted: Caspase-7 | Scleropages Formosus | Fish | Osteoglossiformes |
| NP_001091272.1 | Caspase-7 S Homeolog | Xenopus Laevis | Amphibia | Anura |
| NP_001219.2 | Caspase-8 Isoform A Precursor | Homo Sapiens | Mammalia | Primates |
| NP_001125222.2 | Caspase-8 | Pongo Abelii | Mammalia | Primates |
| NP_001264855.1 | Caspase-8 Isoform 2 | Mus Musculus | Mammalia | Rodentia |
| NP_071613.1 | Caspase-8 | Rattus Norvegicus | Mammalia | Rodentia |
| NP_571585.2 | Caspase-8 | Danio Rerio | Fish | Cypriniformes |
| XP_007663299.1 | Predicted: Caspase-8 | Ornithorhynchus Anatinus | Mammalia | Monotremata |
| NP_001041494.1 | Caspase-8 | Canis Lupus Familiaris | Mammalia | Carnivora |
| NP_989923.1 | Caspase-8 | Gallus Gallus | Birds | Galliformes |
| NP_001026949.2 | Caspase-8 | Sus Scrofa | Mammalia | Artiodactyla |
| NP_001039435.1 | Caspase-8 | Bos Taurus | Mammalia | Artiodactyla |
| XP_017953067.1 | Predicted: Caspase-8 | Xenopus Tropicalis | Amphibia | Anura |
| XP_011367467.1 | Predicted: Caspase-8 | Pteropus Vampyrus | Mammalia | Chiroptera |
| XP_004601328.1 | Predicted: Caspase-8 | Sorex Araneus | Mammalia | Soricomorpha |
| XP_012380773.1 | Predicted: Caspase-8 | Dasypus Novemcinctus | Mammalia | Cingulata |
| NP_001233725.1 | Caspase-8 | Cricetulus Griseus | Mammalia | Rodentia |
| XP_005306309.1 | Predicted: Caspase-8 isoform X1 | Chrysemys Picta Bellii | Reptilia | Testudines |
| XP_006272599.1 | Predicted: Caspase-8 isoform X1 | Alligator Mississippiensis | Reptilia | Crocodilia |
| XP_006272609.1 | Predicted: Caspase-8 | Alligator Mississippiensis | Reptilia | Crocodilia |
| NP_001098258.1 | Caspase-8 | Oryzias Latipes | Fish | Beloniformes |
| XP_009963925.1 | Predicted: Caspase-8 | Tyto Alba | Birds | Strigiformes |
| XP_009807896.1 | Predicted: Caspase-8 | Gavia Stellata | Birds | Gaviiformes |
| XP_009282932.1 | Predicted: Caspase-8 | Aptenodytes Forsteri | Birds | Sphenisciformes |
| XP_009876853.1 | Predicted: Caspase-8 | Apaloderma Vittatum | Birds | Trogoniformes |
| XP_009888484.1 | Predicted: Caspase-8 | Charadrius Vociferus | Birds | Charadriiformes |
| XP_009460551.1 | Predicted: Caspase-8 | Nipponia Nippon | Birds | Pelecaniformes |
| XP_010563026.1 | Predicted: Caspase-8 | Haliaeetus Leucocephalus | Birds | Accipitriformes |
| XP_010404904.1 | Caspase-8 | Corvus Cornix Cornix | Birds | Passeriformes |
| XP_021175080.1 | Caspase-8 | Fundulus Heteroclitus | Fish | Cyprinodontiformes |
| XP_018585033.1 | Predicted: Caspase-8 isoform X1 | Scleropages Formosus | Fish | Osteoglossiformes |
| NP_001187127.1 | Caspase-8 | Ictalurus Punctatus | Fish | Siluriformes |
| NP_001079034.1 | Caspase-8 L Homeolog | Xenopus Laevis | Amphibia | Anura |

| | | | | |
|---|---|---|---|---|
| XP_008929353.1 | Predicted: Caspase-8 | Manacus Vitellinus | Birds | Passeriformes |
| XP_003457507.2 | Predicted: Caspase-8 | Oreochromis Niloticus | Fish | Perciformes |
| XP_021444269.1 | Predicted: Caspase-8 isoform X1 | Oncorhynchus Mykiss | Fish | Salmoniformes |
| XP_022610907.1 | Predicted: Caspase-8 isoform X1 | Seriola Dumerili | Fish | Perciformes |
| NP_116759.2 | Caspase-10 Isoform 1 Preprotein | Homo Sapiens | Mammalia | Primates |
| NP_001127368.1 | Caspase-10 | Pongo Abelii | Mammalia | Primates |
| XP_005640587.1 | Caspase-10 | Canis Lupus Familiaris | Mammalia | Carnivora |
| XP_421936.4 | Predicted: Caspase-10 | Gallus Gallus | Birds | Galliformes |
| NP_001155112.1 | Caspase-10 | Sus Scrofa | Mammalia | Artiodactyla |
| NP_001093436.1 | Caspase-10 | Oryctolagus Cuniculus | Mammalia | Lagomorpha |
| NP_001015715.2 | Caspase-10 | Xenopus Tropicalis | Amphibia | Anura |
| XP_008118735.1 | Predicted: Caspase-10 isoform X1 | Anolis Carolinensis | Reptilia | Squamata |
| XP_004458966.1 | Predicted: Caspase-10 | Dasypus Novemcinctus | Mammalia | Cingulata |
| XP_021591334.1 | Caspase-10 | Ictidomys Tridecemlineatus | Mammalia | Rodentia |
| XP_008175924.1 | Predicted: Caspase-10 isoform X3 | Chrysemys Picta Bellii | Reptilia | Testudines |
| XP_014449789.1 | Predicted: Caspase-10 isoform X1 | Alligator Mississippiensis | Reptilia | Crocodilia |
| XP_004577327.1 | Predicted: Caspase-10 | Ochotona Princeps | Mammalia | Lagomorpha |
| XP_006862096.1 | Predicted: Caspase-10 | Chrysochloris Asiatica | Mammalia | Afrosoricida |
| XP_008144768.1 | Predicted: Caspase-10 | Eptesicus Fuscus | Mammalia | Chiroptera |
| XP_012421961.1 | Predicted: Caspase-10 isoform X1 | Odobenus Rosmarus Divergens | Mammalia | Carnivora |
| XP_015452237.1 | Predicted: Caspase-10 isoform X1 | Pteropus Alecto | Mammalia | Chiroptera |
| XP_014104940.1 | Predicted: Caspase-10 | Pseudopodoces Humilis | Birds | Passeriformes |
| XP_004262912.2 | Predicted: Caspase-10 | Orcinus Orca | Mammalia | Cetacea |
| XP_005484241.1 | Predicted: Caspase-10 | Zonotrichia Albicollis | Birds | Passeriformes |
| XP_014374717.1 | Predicted: Caspase-10 isoform X1 | Alligator Sinensis | Reptilia | Crocodilia |
| XP_007190566.1 | Predicted: Caspase-10 isoform X1 | Balaenoptera Acutorostrata Scammoni | Mammalia | Cetacea |
| XP_009807895.1 | Predicted: Caspase-10 | Gavia Stellata | Birds | Gaviiformes |
| XP_010125639.1 | Predicted: Caspase-10 | Chlamydotis Macqueenii | Birds | Otidiformes |
| XP_008494202.1 | Predicted: Caspase-10 | Calypte Anna | Birds | Apodiformes |
| XP_009876852.1 | Predicted: Caspase-10 | Apaloderma Vittatum | Birds | Trogoniformes |
| XP_009888483.1 | Predicted: Caspase-10 | Charadrius Vociferus | Birds | Charadriiformes |

| | | | | |
|---|---|---|---|---|
| XP_009460550.1 | Predicted: Caspase-10 | Nipponia Nippon | Birds | Pelecaniformes |
| XP_009500813.1 | Predicted: Caspase-10 | Phalacrocorax Carbo | Birds | Suliformes |
| XP_010563039.1 | Predicted: Caspase-10 | Haliaeetus Leucocephalus | Birds | Accipitriformes |
| XP_011574974.1 | Predicted: Caspase-10 | Aquila Chrysaetos Canadensis | Birds | Accipitriformes |
| NP_001081410.1 | Caspase-10 S Homeolog | Xenopus Laevis | Amphibia | Anura |
| NP_001083130.1 | Caspase-10 L Homeolog | Xenopus Laevis | Amphibia | Anura |
| XP_017527989.1 | Predicted: Caspase-10 | Manis Javanica | Mammalia | Pholidota |
| XP_019378751.1 | Predicted: Caspase-10 isoform X1 | Gavialis Gangeticus | Reptilia | Crocodilia |
| NP_001300701.1 | cFLIP | Danio Rerio | Fish | Cypriniformes |
| XP_033025467.1 | cFLIP | Lacerta Agilis | Reptilia | Squamata |
| XP_020667408.1 | cFLIP | Pogona Vitticeps | Reptilia | Squamata |
| XP_009330029.1 | cFLIP | Pygoscelis Adeliae | Birds | Sphenisciformes |
| XP_019409414.1 | cFLIP | Crocodylus Porosus | Reptilia | Crocodilia |
| XP_006022153.1 | cFLIP | Alligator Sinensis | Reptilia | Crocodilia |
| XP_006118011.1 | cFLIP | Pelodiscus Sinensis | Reptilia | Testudines |
| XP_010005644.1 | cFLIP | Chaetura Pelagica | Birds | Apodiformes |
| XP_010123198.1 | cFLIP | Chlamydotis Macqueenii | Birds | Otidiformes |
| XP_009707893.1 | cFLIP | Cariama Cristata | Birds | Cariamiformes |
| XP_030180293.1 | cFLIP | Lynx Pardinus | Mammalia | Carnivora |
| XP_020930360.1 | cFLIP | Sus Scrofa | Mammalia | Artiodactyla |
| XP_022423099.1 | cFLIP | Delphinapterus Leucas | Mammalia | Cetacea |
| XP_026980099.1 | cFLIP | Lagenorhynchus Obliquidens | Mammalia | Cetacea |
| XP_025749261.1 | cFLIP | Callorhinus Ursinus | Mammalia | Carnivora |
| XP_005640588.2 | cFLIP | Canis Lupus Familiaris | Mammalia | Carnivora |
| XP_035161179.1 | cFLIP | Callithrix Jacchus | Mammalia | Primates |
| NP_001120655.1 | cFLIP | Homo Sapiens | Mammalia | Primates |
| NP_001125140.1 | cFLIP | Pongo Abelii | Mammalia | Primates |
| XP_024419473.1 | cFLIP | Desmodus Rotundus | Mammalia | Chiroptera |
| NP_001188445.1 | cFLIP | Oryzias Latipes | Fish | Beloniformes |
| XP_023284372.1 | cFLIP | Seriola Lalandi | Fish | Perciformes |
| XP_020450436.1 | cFLIP | Monopterus Albus | Fish | Synbranchiformes |
| XP_034740679.1 | cFLIP | Etheostoma Cragini | Fish | Perciformes |
| NP_001254595.1 | cFLIP | Gasterosteus Aculeatus | Fish | Gasterosteiformes |
| **AOA 3** | | | | |
| Accension ID | Caspase | Species | Class | Order |

| | | | | |
|---|---|---|---|---|
| NP_004337.2 | Caspase-3 isoform a preprotein | Homo Sapiens | Mammalia | Primates |
| NP_001012435.1 | Caspase-3 | Pan Troglodytes | Mammalia | Primates |
| NP_001271338.1 | Caspase-3 | Mus Musculus | Mammalia | Rodentia |
| NP_037054.1 | Caspase-3 | Rattus Norvegicus | Mammalia | Rodentia |
| NP_571952.1 | Caspase-3 Apoptotsis-related cysteine peptidase a | Danio Rerio | Fish | Cypriniformes |
| NP_001003042.1 | Caspase-3 | Canis Lupus Familiaris | Mammalia | Carnivora |
| NP_001157433.1 | Caspase-3 | Equus Caballus | Mammalia | Perissodactyla |
| NP_990056.1 | Caspase-3 | Gallus Gallus | Birds | Galliformes |
| NP_999296.1 | Caspase-3 | Sus Scrofa | Mammalia | Artiodactyla |
| NP_001071308.1 | Caspase-3 | Bos Taurus | Mammalia | Artiodactyla |
| NP_001075586.1 | Caspase-3 | Oryctolagus Cuniculus | Mammalia | Lagomorpha |
| NP_001120900.1 | Caspase-3 | Xenopus Tropicalis | Amphibia | Anura |
| XP_007905080.1 | Predicted: Caspase-3 | Callorhinchus Milii | Chondrichthyes | Chimaeriformes |
| NP_001230975.1 | Caspase-3 | Cricetulus Griseus | Mammalia | Rodentia |
| XP_006128558.1 | Predicted: Caspase-3 isoform X1 | Pelodiscus Sinensis | Reptilia | Testudines |
| NP_001266895.1 | Caspase-3 | Saimiri Boliviensis | Mammalia | Primates |
| XP_005719459.1 | Predicted: Caspase-3 | Pundamilia Nyererei | Actinopterygii | Cichliformes |
| XP_019336883.1 | Predicted: Caspase-3 | Alligator Mississippiensis | Reptilia | Crocodilia |
| XP_004428794.1 | Predicted: Caspase-3 | Ceratotherium simum simum | Mammalia | Perissodactyla |
| XP_006834493.1 | Predicted: Caspase-3 | Chrysochloris Asiatica | Mammalia | Afrosoricida |
| NP_001098140.1 | Caspase-3 | Oryzias Latipes | Fish | Beloniformes |
| XP_006160500.1 | Predicted: Caspase-3 isoform X2 | Tupaia Chinensis | Mammalia | Scandentia |
| XP_013161017.1 | Predicted: Caspase-3 isoform X1 | Falco Peregrinus | Birds | Falconiformes |
| XP_007054526.1 | Predicted: Caspase-3 | Chelonia Mydas | Reptilia | Testudines |
| XP_014120317.1 | Predicted: Caspase-3 | Zonotrichia Albicollis | Birds | Passeriformes |
| XP_005972660.1 | Predicted: Caspase-3 | Pantholops Hodgsonii | Mammalia | Artiodactyla |
| XP_006026683.1 | Predicted: Caspase-3 | Alligator Sinensis | Reptilia | Crocodilia |
| XP_009099177.1 | Predicted: Caspase-3 isoform X1 | Serinus Canaria | Birds | Passeriformes |
| XP_010281454.1 | Predicted: Caspase-3 | Phaethon Lepturus | Birds | Phaethontiformes |
| XP_009697353.1 | Predicted: Caspase-3 | Cariama Cristata | Birds | Cariamiformes |
| XP_009570651.1 | Predicted: Caspase-3 | Fulmarus Glacialis | Birds | Procellariiformes |
| NP_001290581.1 | Caspase-3 | Esox Lucius | Fish | Esociformes |
| NP_001188010.1 | Caspase-3 | Ictalurus Punctatus | Fish | Siluriformes |
| NP_001081225.1 | Caspase-3 | Xenopus Laevis | Amphibia | Anura |

| | | | | |
|---|---|---|---|---|
| NP_001269823.1 | Caspase-3 | Oreochromis Niloticus | Fish | Perciformes |
| NP_001217.2 | Caspase-6 Isoform Alpha Precursor | Homo Sapiens | Mammalia | Primates |
| NP_033941.3 | Caspase-6 Precursor | Mus Musculus | Mammalia | Rodentia |
| NP_113963.2 | Caspase-6 | Rattus Norvegicus | Mammalia | Rodentia |
| NP_001018333.1 | Caspase-6 | Danio Rerio | Fish | Cypriniformes |
| NP_990057.1 | Caspase-6 | Gallus Gallus | Birds | Galliformes |
| XP_005656604.1 | Predicted: Caspase-6 isoform X1 | Sus Scrofa | Mammalia | Artiodactyla |
| NP_001030496.1 | Caspase-6 | Bos Taurus | Mammalia | Artiodactyla |
| NP_001011068.1 | Caspase-6 | Xenopus Tropicalis | Amphibia | Anura |
| XP_003221840.2 | Predicted: Caspase-6 | Anolis Carolinensis | Reptilia | Squamata |
| XP_014427079.1 | Predicted: Caspase-6 | Pelodiscus Sinensis | Reptilia | Testudines |
| XP_019355646.1 | Predicted: Caspase-6 isoform X1 | Alligator Mississippiensis | Reptilia | Crocodilia |
| XP_014418265.1 | Predicted: Caspase-6 | Camelus Ferus | Mammalia | Artiodactyla |
| XP_004411305.1 | Predicted: Caspase-6 | Odobenus Rosmarus Divergens | Mammalia | Carnivora |
| XP_006918975.1 | Predicted: Caspase-6 isoform X1 | Pteropus Alecto | Mammalia | Chiroptera |
| XP_007054543.1 | Predicted: Caspase-6 | Chelonia Mydas | Reptilia | Testudines |
| XP_014120908.1 | Predicted: Caspase-6 isoform X1 | Zonotrichia Albicollis | Birds | Passeriformes |
| XP_007463292.1 | Predicted: Caspase-6 | Lipotes Vexillifer | Mammalia | Cetacea |
| XP_008315389.1 | Predicted: Caspase-6 | Cynoglossus Semilaevis | Fish | Pleuronectiformes |
| XP_008841740.1 | Predicted: Caspase-6 | Nannospalax Galili | Mammalia | Rodentia |
| XP_008420898.1 | Predicted: Caspase-6 | Poecilia Reticulata | Fish | Cyprinodontiformes |
| XP_009578173.1 | Predicted: Caspase-6 | Fulmarus Glacialis | Birds | Procellariiformes |
| XP_009924247.1 | Predicted: Caspase-6 | Haliaeetus Albicilla | Birds | Accipitriformes |
| XP_017592325.1 | Predicted: Caspase-6 | Corvus Brachyrhynchos | Birds | Passeriformes |
| XP_009930434.1 | Predicted: Caspase-6 | Opisthocomus Hoazin | Birds | Opisthocomiformes |
| XP_010187950.1 | Predicted: Caspase-6 | Mesitornis Unicolor | Birds | Mesitornithiformes |
| XP_019326128.1 | Predicted: Caspase-6 | Aptenodytes Forsteri | Birds | Sphenisciformes |
| XP_012724337.1 | Predicted: Caspase-6 isoform X1 | Fundulus Heteroclitus | Fish | Cyprinodontiformes |
| XP_018425527.1 | Predicted: Caspase-6 | Nanorana Parkeri | Amphibia | Anura |
| XP_019110884.1 | Predicted: Caspase-6 isoform X1 | Larimichthys Crocea | Fish | Perciformes |
| XP_013863217.1 | Predicted: Caspase-6 | Austrofundulus Limnaeus | Fish | Cyprinodontiformes |
| XP_015281944.1 | Predicted: Caspase-6 isoform X1 | Gekko Japonicus | Reptilia | Squamata |
| XP_015666700.1 | Predicted: Caspase-6 | Protobothrops Mucrosquamatus | Reptilia | Squamata |

| | | | | |
|---|---|---|---|---|
| XP_018539675.1 | Predicted: Caspase-6 | Lates Calcarifer | Fish | Perciformes |
| NP_001081406.1 | Caspase-6 L Homeolog | Xenopus Laevis | Amphibia | Anura |
| NP_001117743.1 | Caspase-6 Precursor | Oncorhynchus Mykiss | Fish | Salmoniformes |
| NP_001253985.1 | Caspase-7 Isoform Alpha Precursor | Homo Sapiens | Mammalia | Primates |
| NP_071596.1 | Caspase-7 | Rattus Norvegicus | Mammalia | Rodentia |
| NP_001018443.1 | Caspase-7 | Danio Rerio | Fish | Cypriniformes |
| XP_421764.3 | Predicted: Caspase-7 | Gallus Gallus | Birds | Galliformes |
| NP_001016299.1 | Caspase-7 | Xenopus Tropicalis | Amphibia | Anura |
| XP_008112945.1 | Predicted: Caspase-7 isoform X2 | Anolis Carolinensis | Reptilia | Squamata |
| XP_015091414.1 | Predicted: Caspase-7 | Vicugna Pacos | Mammalia | Artiodactyla |
| XP_004763887.1 | Predicted: Caspase-7 | Mustela Putorius Furo | Mammalia | Carnivora |
| XP_005928237.1 | Predicted: Caspase-7 isoform X1 | Haplochromis Burtoni | Actinopterygii | Cichliformes |
| XP_012410596.1 | Predicted: Caspase-7 | Trichechus Manatus Latirostris | Mammalia | Sirenia |
| XP_013371286.1 | Predicted: Caspase-7 isoform X1 | Chinchilla Lanigera | Mammalia | Rodentia |
| XP_016042302.1 | Predicted: Caspase-7 | Erinaceus Europaeus | Mammalia | Erinaceomorpha |
| XP_005888715.1 | Predicted: Caspase-7 | Bos Mutus | Mammalia | Artiodactyla |
| XP_008142545.1 | Predicted: Caspase-7 isoform X1 | Eptesicus Fuscus | Mammalia | Chiroptera |
| XP_014448895.1 | Predicted: Caspase-7 | Tupaia Chinensis | Mammalia | Scandentia |
| XP_005443932.1 | Predicted: Caspase-7 isoform X1 | Falco Cherrug | Birds | Falconiformes |
| NP_001268771.1 | Caspase-7 | Mesocricetus Auratus | Mammalia | Rodentia |
| XP_006022892.1 | Predicted: Caspase-7 isoform X1 | Alligator Sinensis | Reptilia | Crocodilia |
| XP_009085424.1 | Predicted: Caspase-7 | Serinus Canaria | Birds | Passeriformes |
| XP_008434898.1 | Predicted: Caspase-7 isoform X1 | Poecilia Reticulata | Fish | Cyprinodontiformes |
| XP_008947331.1 | Predicted: Caspase-7 isoform X1 | Merops Nubicus | Birds | Coraciiformes |
| XP_009932919.1 | Predicted: Caspase-7 | Opisthocomus Hoazin | Birds | Opisthocomiformes |
| XP_010180183.1 | Predicted: Caspase-7 isoform X1 | Mesitornis Unicolor | Birds | Mesitornithiformes |
| XP_010010881.1 | Predicted: Caspase-7 | Nestor Notabilis | Birds | Psittaciformes |
| XP_009666939.1 | Predicted: Caspase-7 | Struthio Camelus Australis | Birds | Struthioniformes |
| XP_010895864.1 | Predicted: Caspase-7 | Esox Lucius | Fish | Esociformes |
| XP_020773763.1 | Caspase-7 | Boleophthalmus Pectinirostris | Fish | Perciformes |
| XP_018419981.1 | Predicted: Caspase-7 isoform X1 | Nanorana Parkeri | Amphibia | Anura |

| | | | | |
|---|---|---|---|---|
| XP_010740374.1 | Predicted: Caspase-7 | Larimichthys Crocea | Fish | Perciformes |
| XP_013863024.1 | Predicted: Caspase-7 isoform X1 | Austrofundulus Limnaeus | Fish | Cyprinodontiformes |
| XP_018618189.1 | Predicted: Caspase-7 | Scleropages Formosus | Fish | Osteoglossiformes |
| NP_001081408.1 | Caspase-7 | Xenopus Laevis | Amphibia | Anura |
| NP_001091272.1 | Caspase-7 S Homeolog | Xenopus Laevis | Amphibia | Anura |
| XP_019411759.1 | Predicted: Caspase-7 isoform X1 | Crocodylus Porosus | Reptilia | Crocodilia |
| XP_020635566.1 | Caspase-7 | Pogona Vitticeps | Reptilia | Squamata |
| NP_001219.2 | Caspase-8 Isoform A Precursor | Homo Sapiens | Mammalia | Primates |
| NP_001125222.2 | Caspase-8 | Pongo Abelii | Mammalia | Primates |
| NP_001264855.1 | Caspase-8 Isoform 2 | Mus Musculus | Mammalia | Rodentia |
| NP_071613.1 | Caspase-8 | Rattus Norvegicus | Mammalia | Rodentia |
| NP_571585.2 | Caspase-8 | Danio Rerio | Fish | Cypriniformes |
| NP_001041494.1 | Caspase-8 | Canis Lupus Familiaris | Mammalia | Carnivora |
| NP_989923.1 | Caspase-8 | Gallus Gallus | Birds | Galliformes |
| NP_001026949.2 | Caspase-8 | Sus Scrofa | Mammalia | Artiodactyla |
| NP_001039435.1 | Caspase-8 | Bos Taurus | Mammalia | Artiodactyla |
| XP_017953067.1 | Predicted: Caspase-8 | Xenopus Tropicalis | Amphibia | Anura |
| XP_010711755.1 | Predicted: Caspase-8 | Meleagris Gallopavo | Birds | Galliformes |
| NP_001233725.1 | Caspase-8 | Cricetulus Griseus | Mammalia | Rodentia |
| XP_012778477.1 | Caspase-8 | Maylandia Zebra | Fish | Perciformes |
| XP_005306309.1 | Predicted: Caspase-8 isoform X1 | Chrysemys Picta Bellii | Reptilia | Testudines |
| XP_015214952.1 | Predicted: Caspase-8 | Lepisosteus Oculatus | Holostei | Lepisosteiformes |
| XP_004378277.1 | Predicted: Caspase-8 isoform X1 | Trichechus Manatus Latirostris | Mammalia | Sirenia |
| XP_019830532.1 | Predicted: Caspase-8 | Bos Indicus | Mammalia | Artiodactyla |
| XP_012581231.1 | Predicted: Caspase-8 isoform X1 | Condylura Cristata | Mammalia | Soricomorpha |
| XP_012803201.1 | Predicted: Caspase-8 | Jaculus Jaculus | Mammalia | Rodentia |
| XP_006272599.1 | Predicted: Caspase-8 isoform X1 | Alligator Mississippiensis | Reptilia | Crocodilia |
| XP_006272609.1 | Predicted: Caspase-8 | Alligator Mississippiensis | Reptilia | Crocodilia |
| NP_001098258.1 | Caspase-8 | Oryzias Latipes | Fish | Beloniformes |
| XP_012950190.1 | Caspase-8 isoform X1 | Anas Platyrhynchos | Birds | Anseriformes |
| XP_010017555.1 | Predicted: Caspase-8 | Nestor Notabilis | Birds | Psittaciformes |
| XP_009682548.1 | Predicted: Caspase-8 | Struthio Camelus Australis | Birds | Struthioniformes |
| XP_009894783.1 | Predicted: Caspase-8 | Picoides Pubescens | Birds | Piciformes |
| XP_009876853.1 | Predicted: Caspase-8 | Apaloderma Vittatum | Birds | Trogoniformes |

| | | | | |
|---|---|---|---|---|
| XP_009888484.1 | Predicted: Caspase-8 | Charadrius Vociferus | Birds | Charadriiformes |
| XP_009460551.1 | Predicted: Caspase-8 | Nipponia Nippon | Birds | Pelecaniformes |
| XP_010563026.1 | Predicted: Caspase-8 | Haliaeetus Leucocephalus | Birds | Accipitriformes |
| XP_021175123.1 | Caspase-8 | Fundulus Heteroclitus | Fish | Cyprinodontiformes |
| XP_018585033.1 | Predicted: Caspase-8 isoform X1 | Scleropages Formosus | Fish | Osteoglossiformes |
| NP_001187127.1 | Caspase-8 | Ictalurus Punctatus | Fish | Siluriformes |
| NP_001079034.1 | Caspase-8 L Homeolog | Xenopus Laevis | Amphibia | Anura |
| XP_021444269.1 | Caspase-8 isoform X1 | Oncorhynchus Mykiss | Fish | Salmoniformes |
| NP_116759.2 | Caspase-10 Isoform 1 Preprotein | Homo Sapiens | Mammalia | Primates |
| NP_001127368.1 | Caspase-10 | Pongo Abelii | Mammalia | Primates |
| XP_421936.4 | Predicted: Caspase-10 | Gallus Gallus | Birds | Galliformes |
| NP_001155112.1 | Caspase-10 | Sus Scrofa | Mammalia | Artiodactyla |
| NP_001093436.1 | Caspase-10 | Oryctolagus Cuniculus | Mammalia | Lagomorpha |
| NP_001015715.2 | Caspase-10 | Xenopus Tropicalis | Amphibia | Anura |
| XP_002749670.1 | Predicted: Caspase-10 isoform X1 | Callithrix Jacchus | Mammalia | Primates |
| XP_008118735.1 | Predicted: Caspase-10 isoform X1 | Anolis Carolinensis | Reptilia | Squamata |
| XP_012354911.1 | Predicted: Caspase-10 isoform X1 | Nomascus Leucogenys | Mammalia | Primates |
| XP_008175924.1 | Predicted: Caspase-10 isoform X3 | Chrysemys Picta Bellii | Reptilia | Testudines |
| XP_005049234.1 | Predicted: Caspase-10 | Ficedula Albicollis | Reptilia | Crocodilia |
| XP_014449789.1 | Predicted: Caspase-10 isoform X1 | Alligator Mississippiensis | Reptilia | Crocodilia |
| XP_016042508.1 | Predicted: Caspase-10 | Erinaceus Europaeus | Mammalia | Erinaceomorpha |
| XP_012421961.1 | Predicted: Caspase-10 isoform X1 | Odobenus Rosmarus Divergens | Mammalia | Carnivora |
| XP_015414641.1 | Predicted: Caspase-10 | Myotis Davidii | Mammalia | Chiroptera |
| XP_014445038.1 | Predicted: Caspase-10 isoform X1 | Tupaia Chinensis | Mammalia | Scandentia |
| XP_005573964.1 | Predicted: Caspase-10 | Macaca Fascicularis | Mammalia | Primates |
| XP_005484241.1 | Predicted: Caspase-10 | Zonotrichia Albicollis | Birds | Passeriformes |
| XP_014374717.1 | Predicted: Caspase-10 isoform X1 | Alligator Sinensis | Reptilia | Crocodilia |
| XP_007090058.1 | Predicted: Caspase-10 | Panthera Tigris Altaica | Mammalia | Carnivora |
| XP_009086338.1 | Predicted: Caspase-10 | Serinus Canaria | Birds | Passeriformes |
| XP_009963924.1 | Predicted: Caspase-10 | Tyto Alba | Birds | Strigiformes |
| XP_017583676.1 | Predicted: Caspase-10 isoform X1 | Corvus Brachyrhynchos | Birds | Passeriformes |
| XP_009282933.1 | Predicted: Caspase-10 | Aptenodytes Forsteri | Birds | Sphenisciformes |

| | | | | |
|---|---|---|---|---|
| XP_010165007.1 | Predicted: Caspase-10 | Antrostomus Carolinensis | Birds | Caprimulgiformes |
| XP_010222872.1 | Predicted: Caspase-10 | Tinamus Guttatus | Birds | Tinamiformes |
| XP_013043023.1 | Predicted: Caspase-10 isoform X1 | Anser Cygnoides Domesticus | Birds | Anseriformes |
| XP_014941825.1 | Predicted: Caspase-10 | Acinonyx Jubatus | Mammalia | Carnivora |
| XP_015986581.1 | Predicted: Caspase-10 | Rousettus Aegyptiacus | Mammalia | Chiroptera |
| XP_015489705.1 | Predicted: Caspase-10 | Parus Major | Birds | Passeriformes |
| XP_015723627.1 | Predicted: Caspase-10 isoform X1 | Coturnix Japonica | Birds | Galliformes |
| NP_001081410.1 | Caspase-10 S Homeolog | Xenopus Laevis | Amphibia | Anura |
| NP_001083130.1 | Caspase-10 L Homeolog | Xenopus Laevis | Amphibia | Anura |
| XP_017527989.1 | Predicted: Caspase-10 | Manis Javanica | Mammalia | Pholidota |
| XP_019378751.1 | Predicted: Caspase-10 isoform X1 | Gavialis Gangeticus | Reptilia | Crocodilia |
| NP_001300701.1 | cFLIP | Danio Rerio | Fish | Cypriniformes |
| XP_008118737.1 | cFLIP | Anolis Carolinensis | Reptilia | Squamata |
| XP_033025467.1 | cFLIP | Lacerta Agilis | Reptilia | Squamata |
| XP_025022289.1 | cFLIP | Python Bivittatus | Reptilia | Squamata |
| XP_026709097.1 | cFLIP | Athene Cunicularia | Birds | Strigiformes |
| XP_009479033.1 | cFLIP | Pelecanus Crispus | Birds | Pelecaniformes |
| XP_006022153.1 | cFLIP | Alligator Sinensis | Reptilia | Crocodilia |
| XP_019347600.1 | cFLIP | Alligator Mississippiensis | Reptilia | Crocodilia |
| XP_009976784.1 | cFLIP | Tauraco Erythrolophus | Birds | Musophagiformes |
| XP_009938107.1 | cFLIP | Opisthocomus Hoazin | Birds | Opisthocomiformes |
| XP_025786690.1 | cFLIP | Puma Concolor | Mammalia | Carnivora |
| XP_020930360.1 | cFLIP | Sus Scrofa | Mammalia | Artiodactyla |
| XP_007190580.1 | cFLIP | Balaenoptera Acutorostrata Scammoni | Mammalia | Cetacea |
| XP_026980099.1 | cFLIP | Lagenorhynchus Obliquidens | Mammalia | Cetacea |
| XP_027446588.1 | cFLIP | Zalophus Californianus | Mammalia | Carnivora |
| XP_005640588.2 | cFLIP | Canis Lupus Familiaris | Mammalia | Carnivora |
| XP_019491241.1 | cFLIP | Hipposideros Armiger | Mammalia | Chiroptera |
| XP_035161179.1 | cFLIP | Callithrix Jacchus | Mammalia | Primates |
| NP_001120655.1 | cFLIP | Homo Sapiens | Mammalia | Primates |
| NP_001125140.1 | cFLIP | Pongo Abelii | Mammalia | Primates |
| NP_001188445.1 | cFLIP | Oryzias Latipes | Fish | Beloniformes |
| XP_029385985.1 | cFLIP | Echeneis Naucrates | Fish | Perciformes |
| XP_034417045.1 | cFLIP | Cyclopterus Lumpus | Fish | Scorpaeniformes |
| XP_020506830.2 | cFLIP | Labrus Bergylta | Fish | Perciformes |

| | | | | |
|---|---|---|---|---|
| NP_001254595.1 | cFLIP | Gasterosteus Aculeatus | Fish | Gasterosteiformes |
| **Initiator Lineage** | | | | |
| NP_001219.2 | Caspase-8 Isoform A Precursor | Homo Sapiens | Mammalia | Primates |
| NP_001125222.2 | Caspase-8 | Pongo Abelii | Mammalia | Primates |
| NP_001264855.1 | Caspase-8 Isoform 2 | Mus Musculus | Mammalia | Rodentia |
| NP_071613.1 | Caspase-8 | Rattus Norvegicus | Mammalia | Rodentia |
| NP_571585.2 | Caspase-8 | Danio Rerio | Fish | Cypriniformes |
| NP_001041494.1 | Caspase-8 | Canis Lupus Familiaris | Mammalia | Carnivora |
| NP_989923.1 | Caspase-8 | Gallus Gallus | Birds | Galliformes |
| NP_001026949.2 | Caspase-8 | Sus Scrofa | Mammalia | Artiodactyla |
| NP_001039435.1 | Caspase-8 | Bos Taurus | Mammalia | Artiodactyla |
| XP_017953067.1 | Predicted: Caspase-8 | Xenopus Tropicalis | Amphibia | Anura |
| XP_010711755.1 | Predicted: Caspase-8 | Meleagris Gallopavo | Birds | Galliformes |
| NP_001233725.1 | Caspase-8 | Cricetulus Griseus | Mammalia | Rodentia |
| XP_012778477.1 | Caspase-8 | Maylandia Zebra | Fish | Perciformes |
| XP_005306309.1 | Predicted: Caspase-8 isoform X1 | Chrysemys Picta Bellii | Reptilia | Testudines |
| XP_015214952.1 | Predicted: Caspase-8 | Lepisosteus Oculatus | Holostei | Lepisosteiformes |
| XP_004378277.1 | Predicted: Caspase-8 isoform X1 | Trichechus Manatus Latirostris | Mammalia | Sirenia |
| XP_019830532.1 | Predicted: Caspase-8 | Bos Indicus | Mammalia | Artiodactyla |
| XP_012581231.1 | Predicted: Caspase-8 isoform X1 | Condylura Cristata | Mammalia | Soricomorpha |
| XP_012803201.1 | Predicted: Caspase-8 | Jaculus Jaculus | Mammalia | Rodentia |
| XP_006272599.1 | Predicted: Caspase-8 isoform X1 | Alligator Mississippiensis | Reptilia | Crocodilia |
| XP_006272609.1 | Predicted: Caspase-8 | Alligator Mississippiensis | Reptilia | Crocodilia |
| NP_001098258.1 | Caspase-8 | Oryzias Latipes | Fish | Beloniformes |
| XP_012950190.1 | Caspase-8 isoform X1 | Anas Platyrhynchos | Birds | Anseriformes |
| XP_010017555.1 | Predicted: Caspase-8 | Nestor Notabilis | Birds | Psittaciformes |
| XP_009682548.1 | Predicted: Caspase-8 | Struthio Camelus Australis | Birds | Struthioniformes |
| XP_009894783.1 | Predicted: Caspase-8 | Picoides Pubescens | Birds | Piciformes |
| XP_009876853.1 | Predicted: Caspase-8 | Apaloderma Vittatum | Birds | Trogoniformes |
| XP_009888484.1 | Predicted: Caspase-8 | Charadrius Vociferus | Birds | Charadriiformes |
| XP_009460551.1 | Predicted: Caspase-8 | Nipponia Nippon | Birds | Pelecaniformes |
| XP_010563026.1 | Predicted: Caspase-8 | Haliaeetus Leucocephalus | Birds | Accipitriformes |
| XP_021175123.1 | Caspase-8 | Fundulus Heteroclitus | Fish | Cyprinodontiformes |

| | | | | |
|---|---|---|---|---|
| XP_018585033.1 | Predicted: Caspase-8 isoform X1 | Scleropages Formosus | Fish | Osteoglossiformes |
| NP_001187127.1 | Caspase-8 | Ictalurus Punctatus | Fish | Siluriformes |
| NP_001079034.1 | Caspase-8 L Homeolog | Xenopus Laevis | Amphibia | Anura |
| XP_021444269.1 | Caspase-8 isoform X1 | Oncorhynchus Mykiss | Fish | Salmoniformes |
| NP_116759.2 | Caspase-10 Isoform 1 Preprotein | Homo Sapiens | Mammalia | Primates |
| NP_001127368.1 | Caspase-10 | Pongo Abelii | Mammalia | Primates |
| XP_421936.4 | Predicted: Caspase-10 | Gallus Gallus | Birds | Galliformes |
| NP_001155112.1 | Caspase-10 | Sus Scrofa | Mammalia | Artiodactyla |
| NP_001093436.1 | Caspase-10 | Oryctolagus Cuniculus | Mammalia | Lagomorpha |
| NP_001015715.2 | Caspase-10 | Xenopus Tropicalis | Amphibia | Anura |
| XP_002749670.1 | Predicted: Caspase-10 isoform X1 | Callithrix Jacchus | Mammalia | Primates |
| XP_008118735.1 | Predicted: Caspase-10 isoform X1 | Anolis Carolinensis | Reptilia | Squamata |
| XP_012354911.1 | Predicted: Caspase-10 isoform X1 | Nomascus Leucogenys | Mammalia | Primates |
| XP_008175924.1 | Predicted: Caspase-10 isoform X3 | Chrysemys Picta Bellii | Reptilia | Testudines |
| XP_005049234.1 | Predicted: Caspase-10 | Ficedula Albicollis | Reptilia | Crocodilia |
| XP_014449789.1 | Predicted: Caspase-10 isoform X1 | Alligator Mississippiensis | Reptilia | Crocodilia |
| XP_016042508.1 | Predicted: Caspase-10 | Erinaceus Europaeus | Mammalia | Erinaceomorpha |
| XP_012421961.1 | Predicted: Caspase-10 isoform X1 | Odobenus Rosmarus Divergens | Mammalia | Carnivora |
| XP_015414641.1 | Predicted: Caspase-10 | Myotis Davidii | Mammalia | Chiroptera |
| XP_014445038.1 | Predicted: Caspase-10 isoform X1 | Tupaia Chinensis | Mammalia | Scandentia |
| XP_005573964.1 | Predicted: Caspase-10 | Macaca Fascicularis | Mammalia | Primates |
| XP_005484241.1 | Predicted: Caspase-10 | Zonotrichia Albicollis | Birds | Passeriformes |
| XP_014374717.1 | Predicted: Caspase-10 isoform X1 | Alligator Sinensis | Reptilia | Crocodilia |
| XP_007090058.1 | Predicted: Caspase-10 | Panthera Tigris Altaica | Mammalia | Carnivora |
| XP_009086338.1 | Predicted: Caspase-10 | Serinus Canaria | Birds | Passeriformes |
| XP_009963924.1 | Predicted: Caspase-10 | Tyto Alba | Birds | Strigiformes |
| XP_017583676.1 | Predicted: Caspase-10 isoform X1 | Corvus Brachyrhynchos | Birds | Passeriformes |
| XP_009282933.1 | Predicted: Caspase-10 | Aptenodytes Forsteri | Birds | Sphenisciformes |
| XP_010165007.1 | Predicted: Caspase-10 | Antrostomus Carolinensis | Birds | Caprimulgiformes |
| XP_010222872.1 | Predicted: Caspase-10 | Tinamus Guttatus | Birds | Tinamiformes |
| XP_013043023.1 | Predicted: Caspase-10 isoform X1 | Anser Cygnoides Domesticus | Birds | Anseriformes |

| | | | | |
|---|---|---|---|---|
| XP_014941825.1 | Predicted: Caspase-10 | Acinonyx Jubatus | Mammalia | Carnivora |
| XP_015986581.1 | Predicted: Caspase-10 | Rousettus Aegyptiacus | Mammalia | Chiroptera |
| XP_015489705.1 | Predicted: Caspase-10 | Parus Major | Birds | Passeriformes |
| XP_015723627.1 | Predicted: Caspase-10 isoform X1 | Coturnix Japonica | Birds | Galliformes |
| NP_001081410.1 | Caspase-10 S Homeolog | Xenopus Laevis | Amphibia | Anura |
| NP_001083130.1 | Caspase-10 L Homeolog | Xenopus Laevis | Amphibia | Anura |
| XP_017527989.1 | Predicted: Caspase-10 | Manis Javanica | Mammalia | Pholidota |
| XP_019378751.1 | Predicted: Caspase-10 isoform X1 | Gavialis Gangeticus | Reptilia | Crocodilia |
| NP_001300701.1 | cFLIP | Danio Rerio | Fish | Cypriniformes |
| XP_008118737.1 | cFLIP | Anolis Carolinensis | Reptilia | Squamata |
| XP_033025467.1 | cFLIP | Lacerta Agilis | Reptilia | Squamata |
| XP_025022289.1 | cFLIP | Python Bivittatus | Reptilia | Squamata |
| XP_026709097.1 | cFLIP | Athene Cunicularia | Birds | Strigiformes |
| XP_009479033.1 | cFLIP | Pelecanus Crispus | Birds | Pelecaniformes |
| XP_006022153.1 | cFLIP | Alligator Sinensis | Reptilia | Crocodilia |
| XP_019347600.1 | cFLIP | Alligator Mississippiensis | Reptilia | Crocodilia |
| XP_009976784.1 | cFLIP | Tauraco Erythrolophus | Birds | Musophagiformes |
| XP_009938107.1 | cFLIP | Opisthocomus Hoazin | Birds | Opisthocomiformes |
| XP_025786690.1 | cFLIP | Puma Concolor | Mammalia | Carnivora |
| XP_020930360.1 | cFLIP | Sus Scrofa | Mammalia | Artiodactyla |
| XP_007190580.1 | cFLIP | Balaenoptera Acutorostrata Scammoni | Mammalia | Cetacea |
| XP_026980099.1 | cFLIP | Lagenorhynchus Obliquidens | Mammalia | Cetacea |
| XP_027446588.1 | cFLIP | Zalophus Californianus | Mammalia | Carnivora |
| XP_005640588.2 | cFLIP | Canis Lupus Familiaris | Mammalia | Carnivora |
| XP_019491241.1 | cFLIP | Hipposideros Armiger | Mammalia | Chiroptera |
| XP_035161179.1 | cFLIP | Callithrix Jacchus | Mammalia | Primates |
| NP_001120655.1 | cFLIP | Homo Sapiens | Mammalia | Primates |
| NP_001125140.1 | cFLIP | Pongo Abelii | Mammalia | Primates |
| NP_001188445.1 | cFLIP | Oryzias Latipes | Fish | Beloniformes |
| XP_029385985.1 | cFLIP | Echeneis Naucrates | Fish | Perciformes |
| XP_034417045.1 | cFLIP | Cyclopterus Lumpus | Fish | Scorpaeniformes |
| XP_020506830.2 | cFLIP | Labrus Bergylta | Fish | Perciformes |
| NP_001254595.1 | cFLIP | Gasterosteus Aculeatus | Fish | Gasterosteiformes |
